# Inhibitory potential of autologous neutralizing antibodies sets quantitative limits on the rebound-competent HIV-1 reservoir

**DOI:** 10.64898/2025.12.07.692769

**Authors:** Mauro A. Garcia, Anna Farrell-Sherman, Beril Aydin, Junlin Zhuo, Emily J. Fray, Anna M. Zinsser, Dylan H. Westfall, Kirsten Sowers, Haoyue Li, Brianna M. Lopez, Anthony Abeyta-Lopez, Tifany Chu, Donald Lubbeck, Moonki Chae, Niklas Bachmann, Joseph Varriale, Jun Lai, Francesco R. Simonetti, Rebecca Hoh, Thomas Dalhuisen, Steven G. Deeks, Michael J. Peluso, Janet D. Siliciano, Lillian B. Cohn, Robert F. Siliciano

## Abstract

HIV-1 cure requires preventing viral rebound after treatment interruption, but quantitative criteria defining the rebound-competent reservoir are lacking. We studied individuals undergoing observational treatment interruption without confounding interventions to identify virologic and immunologic determinants of rebound. In 9 of 13 participants, rebound viruses were genetically identical or similar to proviruses in circulating resting CD4^+^ T-cells. We found no evidence of recombination among rebound sequences. Instead, resistance to autologous neutralizing antibodies (aNAbs) was a critical determinant of viral rebound. Increased suppression of viral outgrowth by contemporaneous IgG isolated from plasma was correlated with longer time to rebound. Using inhibitory potential (*IP*), the log reduction in single-round infection at physiologic IgG concentrations, we defined quantitative limits governing rebound-competency with respect to contemporaneous aNAbs. Contemporaneous IgG antibodies inhibited different reservoir variants with a wide range of *IP* values (0.4-8.2 logs), whereas rebound viruses were minimally inhibited (0.5-2.8 logs), indicating that inhibition by even up to 2.8 logs (631-fold) cannot prevent rebound. Longitudinal analyses revealed that waning aNAb potency over time on ART allows previously neutralized variants to gain rebound potential, consistent with the finding that rebound can come from variants deposited in the reservoir at different pre-ART time points. Thus, rebound competency is a dynamic, immune-governed property defined by quantitative immunologic constraints, including those exerted by aNAbs.

**SIGNIFICANCE STATEMENT:** Preventing viral rebound after treatment interruption is the goal of HIV-1 cure research, but the latent proviruses responsible remain undefined. Although rebound is initiated in lymphoid tissues, we found rebound viruses are genetically similar to proviruses in circulating resting CD4^+^ T-cells. Rebound is not explained by recombination and is not solely from proviruses seeded at treatment initiation. Instead, rebound potential is governed by autologous neutralizing antibodies (aNAbs). We define a quantitative threshold of aNAb-mediated inhibition identifying reservoir variants with rebound potential. During treatment, waning aNAb levels allow previously neutralized variants to become rebound-competent. Thus rebound-competency is not a static property, but a dynamic immune-governed feature. Durable aNAb responses against all rebound-competent reservoir variants may be required for functional HIV-1 cure.

## INTRODUCTION

A major barrier to HIV-1 cure is the reservoir of latent proviruses that persist in resting CD4^+^ T cells^1–3^ and rekindling viral replication following interruption of antiretroviral therapy (ART)^4–8^. One approach to HIV-1 cure is elimination of this latent reservoir through the shock-and-kill strategy^9^. Another approach, termed functional cure, involves enhancing viral immunity to prevent post-ART rebound. Recent investigations of rebound have focused on the cell type, differentiation state, proviral integration site, and anatomical location of the cells initiating viral rebound^5,6,10–17^. Rebound competency has been considered an intrinsic, static feature of a fixed subset of persistent proviruses, namely those lacking common fatal defects^18^. Yet, this important subset has not been definitively defined by functional or quantitative criteria.

A critical step toward defining rebound competency is understanding how reservoir proviruses relate to the viruses that actually rebound following interruption of ART. During untreated infection, HIV-1 evolves rapidly, accumulating substantial genetic diversity relative to the transmitted/founder sequence and continually seeding the reservoir with diverse variants^19–23^. In contrast, rebound viremia is often oligoclonal, and, surprisingly, rebound viruses are often phylogenetically distinct even from proviruses in dominant clones of infected CD4^+^ T cells detected using viral outgrowth assays^11,24–27^. This finding led to the hypothesis that recombination of viral variants is involved in rebound^11,25–27^. In addition, recent studies show a modest overrepresentation in the reservoir of variants seeded close to the time of ART initiation, suggesting that these variants may be the source of rebound^21–23,28^.

We recently hypothesized that the host immune response is a critical factor in determining which variants cause rebound^29^. During untreated infection, autologous neutralizing antibodies (aNAbs) exert strong selective pressure on HIV-1, driving the emergence of escape variants and shaping the composition of the circulating plasma virus population^30,31^. Unlike broadly neutralizing antibodies (bNAbs)^32–34^, which have activity against HIV-1 variants from many different people with HIV (PWH), aNAbs are highly specific for HIV-1 envelope (Env) variants present in a given PWH. We have shown that the reservoir preserves a genetically diverse archive of proviruses with varying susceptibility to contemporaneous aNAbs. Only a subset of these reservoir viruses can replicate exponentially *ex vivo* in the presence of low concentrations of contemporaneous, autologous IgG^1^, a finding that has generated additional interest in the role of aNAbs^31–35^. Following ART initiation, viremia is suppressed to undetectable levels due to the short half-lives of most productively infected cells^40–42^, thus diminishing viral antigen exposure and potentially reducing the immune stimulation necessary to generate and sustain antibody production to viral variants. Esmaeilzadeh et al. have shown that aNAbs can emerge after early ART initiation and affect viral rebound^35^. Together, these studies raise questions about the potential of aNAbs in HIV-1 cure strategies.

To explore these questions, we analyzed longitudinal samples from participants undergoing analytic treatment interruption (ATI), integrating intensive HIV-1 *env* sequencing across pre-ART plasma, proviral DNA, outgrowth-derived viruses, and rebound plasma virus with neutralization assays using autologous IgG obtained before and during ART suppression. This combined genetic and functional approach enabled us to explore how cellular compartments, reservoir seeding times, and recombination are related to rebound-competency. Most importantly, we demonstrated a link between aNAb activity and time to viral rebound, defined a minimum quantitative threshold for aNAb activity required to prevent rebound, and demonstrated that temporal changes in aNAb activity reshape the rebound-competent HIV-1 reservoir, revealing a dynamic interplay between reservoir proviruses and the immune response.

## RESULTS

### Study participant characteristics and sampling

The Researching Early Biomarkers of Unsuppressed HIV Dynamics (REBOUND) study (Clinical Trials.gov NCT04359186) enrolled 20 people with HIV-1 (PWH) who maintained suppression of viremia on ART (median=9.5 years) and consented to a supervised ATI. This is one of the few ATI studies without cure-directed interventions, enabling direct evaluation of how aNAbs shape rebound in the absence of confounding effects from interventions such as therapeutic vaccines or broadly neutralizing antibody (bNAb) infusions^8,24,25,43^.

Fourteen REBOUND participants who had undergone baseline leukapheresis to provide sufficient resting CD4^+^ T cells for multiple reservoir assays were included. Resting CD4^+^ T cells were purified from >500 million PBMCs, permitting proviral DNA sequencing, modified quantitative viral outgrowth assays (mQVOAs), and intact proviral DNA assays (IPDA). Leukapheresis was performed a median of 9 weeks prior to ATI (range=3-30 weeks), at which time autologous IgG was also purified for neutralization assays.

The cohort included 9 standard progressors (SPs) who initiated ART during chronic infection, two viremic controllers (VCs) who maintained plasma HIV-1 RNA ≤2,000 copies/mL with stable CD4^+^ counts before initiation of ART, and two elite controllers (ECs) maintained HIV-1 RNA a below the clinical limit of detection (30 copies/mL) without treatment for years, with preserved CD4^+^ levels, before eventual initiation of ART based on current treatment guidelines.^44^ ECs possessed HLA class I alleles previously linked to durable viral control^45–49^; HLA-B*27:05 in EC107 and HLA-B*14:02 in EC106. Neither possess the CCR5Δ32 deletion. The final participant, SP206, initiated ART during hyperacute infection (HAI), 10 days after exposure, and did not develop detectable HIV-1-specific antibodies^50^, providing a rare opportunity to evaluate viral rebound in the absence of measurable aNAb responses. Detailed clinical data are provided in Extended Data Table 1 and visualized in Extended Data Fig. 1.

Participants experienced viral rebound between 7 and 81 days (median= 19 days) following ART interruption. Longitudinal trajectories of plasma HIV-1 RNA, CD4^+^ counts, ART exposure, and sampling for representative SPs, VCs, and ECs are shown in Fig. 1 (and for all participants in Extended Data Fig.1). ART was restarted in SPs once plasma HIV-1 RNA reached ≥30 copies/mL. Due to the ∼1 week processing time for viral load assays, maximum plasma HIV-1 RNA reached in SPs ranged from 268 to 85,686 copies/mL. VC and EC voluntarily restarted ART at the discretion of the individual and their physicians. All participants were on Integrase Strand Transfer Inhibitor (INSTI)-based regimens at the time of ATI; none were on Non-nucleoside Reverse Transcriptase Inhibitor (NNRTI)-based regimens, thus avoiding potential biases from prolonged NNRTI pharmacokinetics (“drug tail” effects), previously linked to delayed rebound^8,51^.

**Figure 1.**
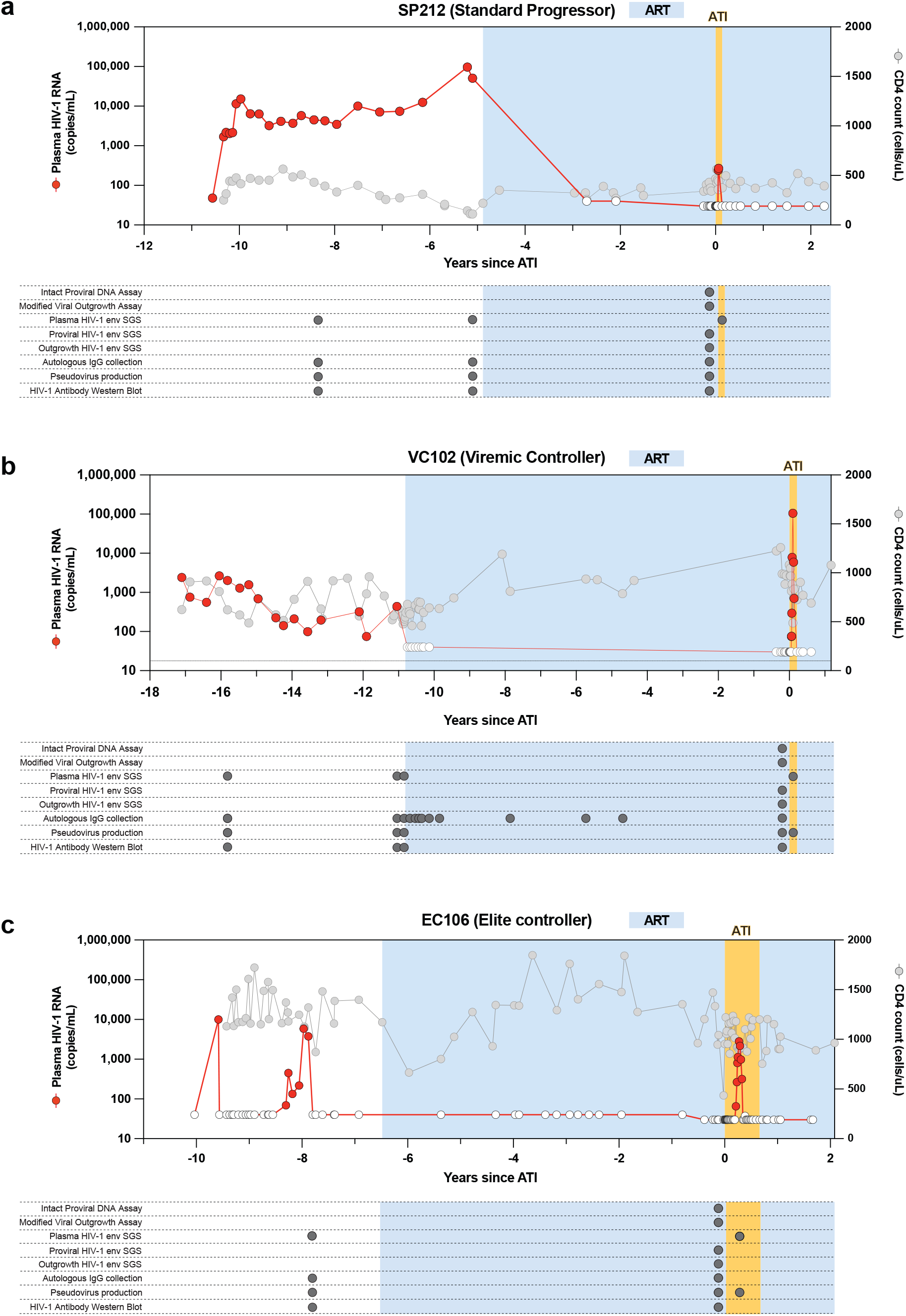
Study participant characteristics, ART interruption, and sampling overview. Longitudinal clinical trajectories for representative participants from each phenotype: (a) standard progressor, (b) viremic controller, and (c) elite controller are shown including plasma HIV-1 RNA (left y-axis, red), CD4^+^ T-cell counts (right y-axis, gray), ART exposure (blue shading), and timepoints for biological sampling for *env* sequencing, antibody purification, and pseudovirus production are indicated with gray circles in the horizontal time track. Gold shading indicates the analytic treatment interruption (ATI) period. Detailed clinical parameters, including ART regimen, duration, and CD4 nadir, are provided in Supplementary Table 1.

To characterize rebound viremia, a total of 1,090 rebound plasma *env* sequences were recovered across participants (median=41; range=4-323), providing broad coverage of viruses present during the initial phase of rebound following ART interruption. To assess the depth of rebound sampling, we performed a power analysis, as previously described^52^. For 9 of 14 participants, there was >90% probability that all rebound lineages comprising ≥10% of the plasma virus population were sequenced. Even for the least sampled participants, there was ≥95% probability of capturing dominant lineages comprising ≥25-50% of the initial rebounding population (Extended Data Table 2).

For comparison to the rebound sequences, we generated HIV-1 env sequences from proviral DNA and outgrowth assay-derived viruses from resting CD4^+^ T cells using single-genome sequencing (SGS). We also sequenced *env* genes from pre-ART plasma virus. Historical pre-ART plasma samples were available for 8 of 14 participants (n=4 SP, n=2 VC, n=2 EC), allowing for characterization of viruses circulating in plasma prior to ART initiation and comparison with reservoir and rebound sequences. Across all participants, a total of 2,260 HIV-1 *env* sequences from pre-ART plasma virus, proviruses in CD4^+^ T cells, outgrowth assay-derived viruses, and rebound isolates were included in our analyses (median=145 per participant, range=37-357) (Extended Data Table 2). A maximum-likelihood phylogenetic tree of unique sequences from each participant, rooted to HXB2, confirmed independent viral evolution in each participant and demonstrated striking inter-participant differences in sequence diversification (Extended Data Fig. 2). Thus, the REBOUND ATI study provided a rigorously controlled and intensively sampled cohort to investigate the virologic and immunologic determinants of HIV-1 rebound in the absence of confounding immunologic or pharmacologic interventions.

### Reservoir variants with reduced susceptibility to pre-ATI autologous antibodies seed initial viral rebound

Autologous neutralizing antibodies (aNAbs) shape viral evolution during untreated infection^30,31^, raising the question of whether they also influence which reservoir variants reemerge during rebound. To assess the extent to which aNAbs select which reservoir viruses rebound during ATI, polyclonal IgG was purified from plasma collected prior to ATI. The presence of HIV-1 Env-specific antibody responses was assessed by Western blot (Extended Data Fig. 3a). Purified IgG was then tested for its ability to neutralize pseudoviruses expressing Env from rebound viruses and from reservoir viruses identified by single-genome sequencing (SGS) of proviral DNA and viral outgrowth assays derived from resting CD4^+^ T cells collected a median of 9 weeks prior to ATI. Neutralization potency was quantified by the half-maximal inhibitory concentration (*IC*_*50*_), the IgG concentration required for 50% inhibition, and by inhibitory potential (*IP*), a previously described^53–55^ metric which integrates both the *IC*_*50*_ and the slope (m) of the dose-response curve to estimate the degree of antibody-mediated inhibition at physiologic antibody concentrations (∼10 mg/mL). *IP*_*10mg/mL*_ is the expected logarithmic reduction in single round infection events at a physiologic concentration of IgG.

Reservoir isolates, including replication-competent isolates from outgrowth assays, exhibited substantial heterogeneity in sensitivity to contemporaneous autologous IgG, whereas rebound isolates were uniformly resistant. Representative dose-response curves from SP209 illustrate the extremes of aNAb sensitivity (Fig. 2a, 2b). A latent reservoir isolate (pseudovirus 209.PRV.5) was potently neutralized by pre-ATI IgG (*IC*_*50*_ = 9.47 µg/mL), with a classic sigmoidal dose-response curve in which infection decreased with increasing antibody concentration (Fig. 2a). In contrast, a rebound isolate from the same participant (pseudovirus 209.REB.5) exhibited markedly reduced sensitivity to pre-ATI IgG (Fig. 2b). Inhibition of infection was observed at the highest antibody concentration tested but did not reach 50% inhibition. As discussed below, the median effect equation^53–55^ can be used to estimate the *IC*_*50*_. For this isolate, the *IC*_*50*_ was 315 µg/mL, requiring 33-fold (1.52 log_10_) higher autologous IgG concentrations to achieve 50% inhibition than for 209.PRV.5. These isolates differ at 55 amino acid positions within the env gene, with a notable concentration of differences in V1 (7) and V4 (11) likely resulting from aNAb pressure.

**Figure 2.**
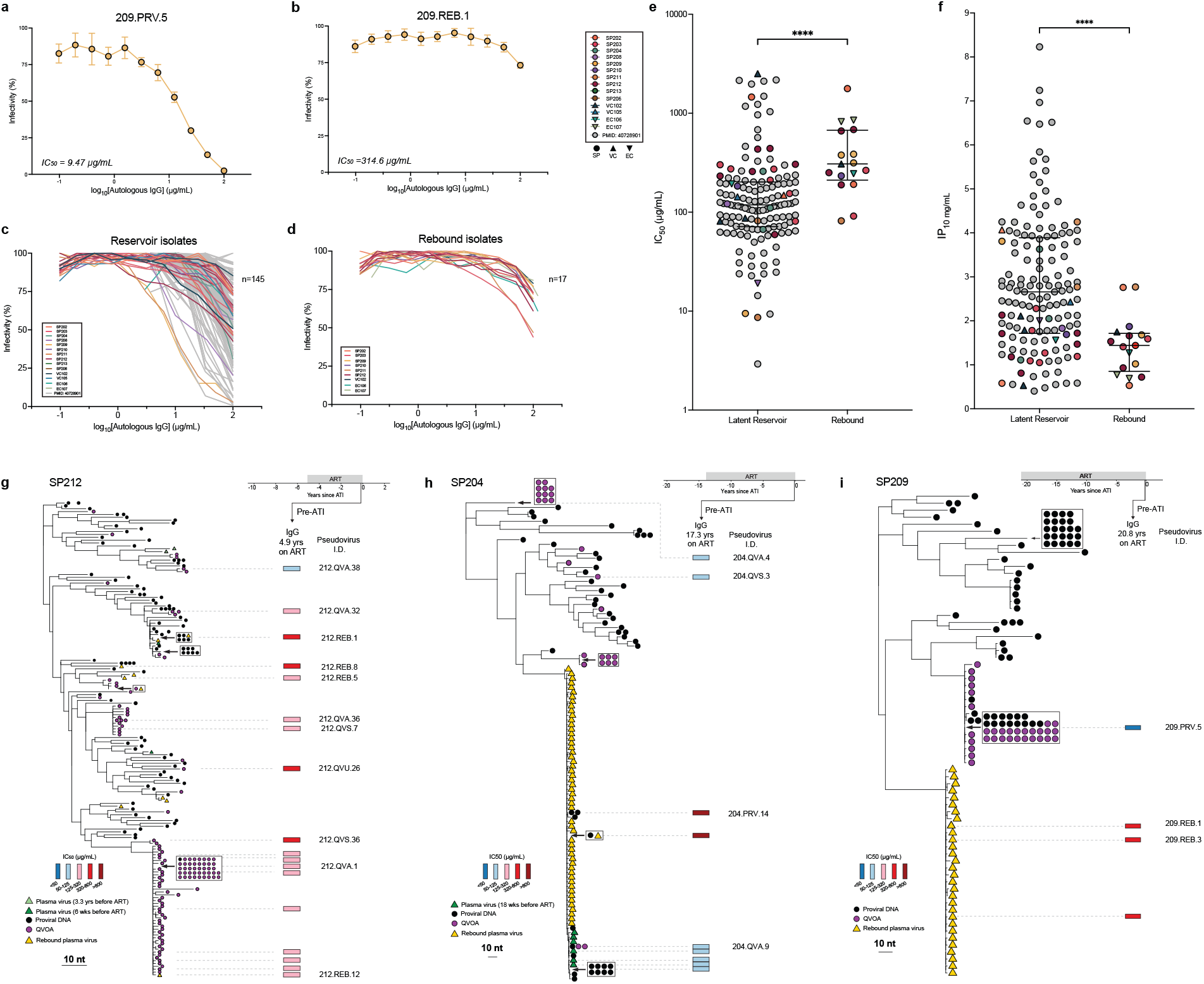
Reservoir variants with reduced susceptibility to pre-ATI autologous antibodies seed initial viral rebound. (a-b) Representative neutralization curves from participant SP209 tested against autologous IgG purified about 2 months prior to the treatment interruption trial; bars indicate mean±SEM. (a) A potently neutralized proviral reservoir isolate, 209.PRV.5, and (b) a minimally inhibited rebound isolate 209.REB.1. (c) Dose-response curves for aNAb-mediated inhibition of infection by reservoir-derived pseudoviruses (n=145). Pseudoviruses from the REBOUND cohort (n=51) were tested against autologous IgG isolated before the ATI (colored curves). For context, reported curves for a broader set of reservoir isolates from PWH on stable ART tested against contemporaneous IgG are included (gray curves). (d) Dose-response curves for aNAb-mediated inhibition of infection by initial rebound pseudoviruses from the REBOUND cohort (n= 17) were tested against autologous IgG isolated before the ATI. (e) Comparison of *IC*_*50*_ values between reservoir viruses from Fig. 2c and rebound viruses from Fig. 2d. Values for reservoir pseudoviruses from the ATI cohort (colored symbols) are within range seen for the reference cohorts (gray symbols) and on average lower than *IC*_*50*_ values for the rebound isolates. The difference between the distributions of reservoir and rebound log_10_-transformed *IC*_*50*_ values was significant as assessed by Mann-Whitney test (p < 0.0001). Bars indicate the median and IQR. (f) *IP*_*10mg/mL*_ values for the isolates in Fig. 2e calculated using the median effect equation. Significance was assessed by Mann-Whitney test (p < 0.0001). Bars indicate the median and IQR. (g-i) Maximum-likelihood *env* phylogenies (bootstrap replicates: n=1,000) for representative participants (SP212, SP204, SP209) with neutralization sensitivity mapped in relation to a blue (more sensitive) to red (more resistant) color scale; rebound viruses cluster on branches minimally inhibited by autologous IgG. Pre-ART plasma virus sequences (green triangles), proviral DNA (black circles), QVOA-derived isolates (purple circles), and rebound plasma viruses (yellow triangles) are represented on midpoint rooted *env* phylogenies. Horizontal timelines indicate autologous IgG sampling, with arrows denoting the specific timepoints used for each neutralization assay in relation to time on ART; continuous suppressive ART before the ATI is indicated by the dark gray shaded region on the horizontal timeline. Pseudovirus IDs are displayed to the right of the neutralization heatmaps. Viral isolates with distinct nucleotide sequences but 100% identical amino acid sequences share identical *IC*_*50*_ values, reflecting equivalent neutralization phenotypes.

We then visualized dose-response curves for unique reservoir isolates from the REBOUND cohort (n=28), together with additional reservoir isolates (n=117) tested from 23 other people with HIV-1 on suppressive ART^56^ to provide a more comprehensive view of the full range of aNAb sensitivities among viruses archived in the latent reservoir during suppressive ART (Fig. 2c). These additional isolates were obtained from individuals sampled during suppressive ART and do not have paired rebound viruses available for comparison. These individuals had durations of time on ART comparable to those of the REBOUND ATI cohort overall (mean 15.3 vs. 14.9 years; median 16.6 vs. 12.3 years, respectively). Rebound viruses are limited for most people with HIV-1 because treatment interruption is generally avoided outside carefully monitored ATI trials, many of which include immunologic or antibody-based interventions that can confound rebound phenotypes. Therefore, reservoir isolates (n=145) were tested in our laboratory using identical experimental conditions for proviral and outgrowth *env* SGS, cloning and generation of pseudoviruses, purification of contemporaneous autologous IgG, and TZM-bl neutralization assays^56^. Reservoir isolates (n=145) displayed a wide range of neutralization sensitivity to contemporaneous, autologous IgG with some isolates exhibiting classic sigmoidal inhibition curves and others showing minimal downward deflection even at the highest antibody concentration tested *in vitro* (Fig. 2c). Curves for unique initial rebound isolates from the REBOUND cohort (n =17) showed downward deflection primarily at antibody concentrations approaching 100 µg/mL (Fig. 2d).

To calculate *IC*_*50*_ values for all isolates, including those for which infectivity was not reduced to below 50%, even at the highest antibody concentration tested *in vitro*, we linearized dose-response curves using a form of the median-effect equation:

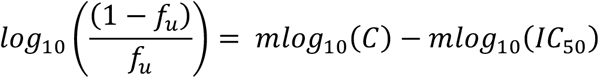

where *f*_*u*_ is the fraction of single round infection events unaffected by the antibody, *m* is the dose-response curve slope or Hill coefficient, and *C* is the antibody concentration in μg/mL, as previously described^53–55^. We then asked whether rebound isolates occupy the same distribution of neutralization phenotypes as reservoir isolates, or whether they represent a shifted subset characterized by reduced aNAb sensitivity. To address this question, we compared log-transformed *IC*_*50*_ values between independent groups of reservoir and rebound isolates using a two-sample Mann-Whitney test. This nonparametric, rank-based method evaluates whether two independent samples differ in distribution; this method is appropriate for continuous data with unequal sample sizes, unpaired observations, and non-normal distributions^57,58^. The null hypothesis (H_0_) being tested was that the distribution of log-transformed *IC*_*50*_ values is identical for both reservoir and rebound isolates. Rejection of this null hypothesis indicates that rebound viruses are a phenotypically distinguished subset within the latent reservoir.

Reservoir isolates exhibited a broad of sensitivity to contemporaneous, autologous IgG (*IC*_*50*_=2.93-2512 µg/mL), whereas rebound isolates uniformly exhibited *IC*_*50*_ values >80 µg/mL (Fig. 2e). Rebound isolates had a median *IC*_*50*_ of 308 µg/mL (IQR: 211–673), significantly higher than reservoir variants (median 119 µg/mL; IQR: 71–202; p < 0.0001, Mann-Whitney test on log-transformed *IC*_*50*_ values). Therefore, we reject the null hypothesis and conclude that rebound isolates are not drawn from the same phenotypic distribution as reservoir isolates. Importantly, if the analysis is restricted to reservoir isolates from the more limited number of PWH for whom rebound isolates are available, the difference in *IC*_*50*_ values is also significant (p= 0.0095, Mann-Whitney test, Extended Data Fig. 3b). These data indicate that this distributional shift does not depend on cross-cohort effects and supports selective replication of autologous neutralization-resistant reservoir variants during initial viral rebound, consistent with previous findings^35^.

Physiologic IgG concentrations are orders of magnitude higher than the concentrations used in *in vitro* neutralization assays. To obtain a better idea of the inhibitory activity of aNAbs at an average physiologic IgG concentration (10 mg/mL), we calculated *IP*_*10mg/mL*_ values for each isolate. This value represents the expected number of logs of inhibition of single round infection events at that concentration. *IP*_*10mg/mL*_ values were determined using the equation^53–55^:

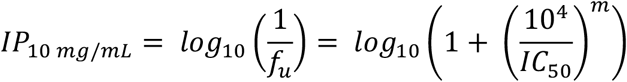

Importantly, by incorporating both the *IC*_*50*_ and the slope of the dose-response curve, our analysis provides a more complete estimate of inhibition by aNAbs at physiological concentrations. Previous studies have shown that the slope parameter is an extremely important determinant of antiviral activity of antiretroviral drugs and broadly neutralizing antibodies^53–55,59^.

*IP*_*10mg/mL*_ values for reservoir isolates varied over an extremely wide range of almost 8 logs (0.4 to 8.2 logs), whereas *IP*_*10mg/mL*_ values for rebound isolates were ≤ 2.77 logs. (Fig. 2f). Thus, initial rebound came from the subset of reservoir variants for which pre-ATI IgG had reduced inhibitory potential. The median *IP*_*10mg/mL*_ value for rebound viruses was 1.44 logs (IQR: 0.85-1.72), significantly lower (Mann-Whitney test, p<0.0001, Fig. 2f) than reservoir viruses with a median *IP*_*10mg/mL*_ value of 2.66 logs (IQR: 1.72-3.89). These data empirically define a minimum *IP*_*10mg/mL*_ threshold of ∼2.8 log below which aNAbs cannot prevent viral rebound. It is possible that this minimum value could increase if other initial rebound isolates with higher *IP*_*10mg/mL*_ values are identified.

To contextualize these functional phenotypes in a phylogenetic framework, neutralization sensitivity was mapped alongside maximum-likelihood HIV-1 *env* phylogenies. In representative participants (SP212, SP204, SP209), rebound viruses consistently localized to reservoir lineages that were highly resistant to aNAbs at the time of treatment interruption, whereas potently neutralized reservoir lineages did not contribute to viral rebound (Fig. 2g-i). Together, these functional analyses establish that HIV-1 rebound originates from reservoir lineages with reduced susceptibility to contemporaneous autologous IgG. These results provide evidence that viral rebound is not an entirely stochastic process, but rather reflects the selection of reactivated reservoir variants that cannot be sufficiently neutralized by aNAbs at the time of ART interruption.

### Pre-ATI autologous IgG activity correlates with in vivo viral rebound timing and kinetics

We next evaluated whether the potency and breadth of pre-ATI IgG neutralizing activity against autologous viruses directly influenced *in vivo* rebound timing and kinetics. We first tested whether pre-ATI IgG could suppress exponential viral outgrowth from the inducible, replication-competent reservoir. Modified quantitative viral outgrowth assays (mQVOAs) were performed in which resting CD4^+^ T cells purified from samples collected a median of 9 weeks prior to ATI (range=3-30 weeks), were reactivated *ex vivo* in the presence of no IgG, HIV-negative donor IgG (50 µg/mL), or contemporaneous pre-ATI autologous IgG (50 µg/mL), as previously described^29^.

This analysis revealed participant-specific differences in functional neutralizing antibody activity. For SP209, pre-ATI autologous IgG had minimal impact on *ex vivo* viral outgrowth (Fig. 3a). The frequency of proviruses capable of giving rise to viral outgrowth in the presence of control IgG was 9.82 infection units per million (IUPM) resting CD4^+^ T cells (95% CI: 7.74-12.5 IUPM), and in the presence of autologous IgG the frequency was 10.5 IUPM (95% CI: 8.28-13.4). In contrast, for SP208, pre-ATI autologous IgG potently suppressed *ex vivo* viral outgrowth, reducing IUPM values from 8.39 (95% CI: 6.63-10.6) to 0.39 (95% CI: 0.19-0.78). This represents a 95.4% reduction of viral outgrowth by aNAbs (Fig. 3b). EC106 exhibited minimal viral outgrowth, with only 1 of 321 wells scoring positive for viral outgrowth (Fig. 3c); this is consistent with the low frequency of inducible proviruses in elite controllers^30-32^. Robust viral outgrowth sufficient to quantify reductions in IUPM was generally not observed for ECs or VCs, and thus analyses of antibody-mediated suppression were restricted to SPs. We calculated the IUPM and percent reduction in IUPM mediated by aNAbs compared to control IgG in the modified QVOA (see Methods). aNAb-mediated suppression of viral outgrowth ranged from no measurable suppression to a 95.4% reduction in IUPM (Fig. 3d), consistent with previous findings^29^. A full summary of mQVOA outgrowth, HIV-1 p24 ELISA readouts, and IUPM values with 95% confidence intervals are provided in Extended Data Fig. 4 for all participants.

**Figure 3.**
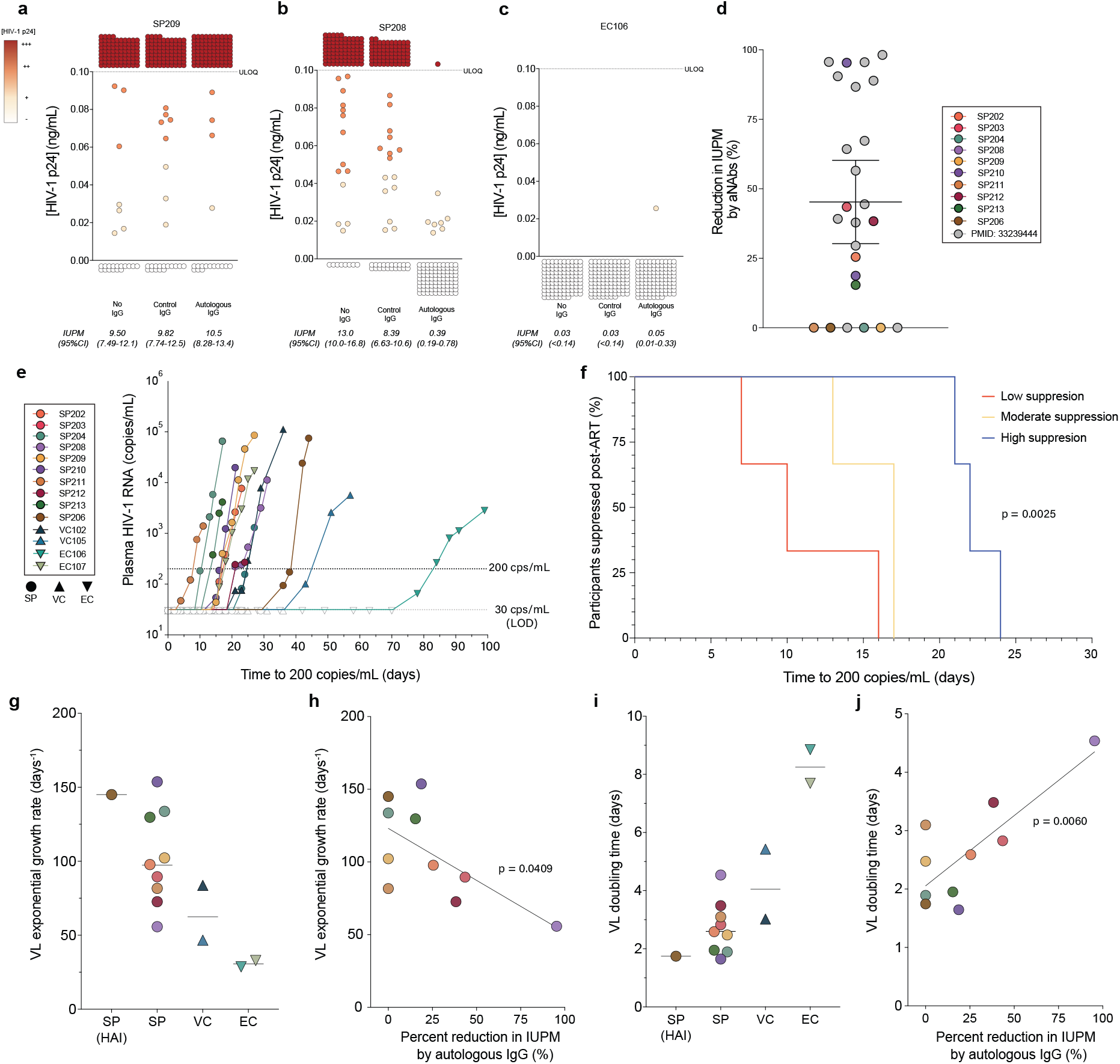
Pre-ATI autologous antibody activity correlate with *in vivo* rebound timing and kinetics. (a-c) Modified quantitative viral outgrowth assays (mQVOAs) from representative participants showing variable suppression of *ex vivo* viral outgrowth by pre-ATI autologous IgG (50 µg/mL) compared to control arms with no IgG, or IgG (50 µg/mL) purified from HIV-negative donors. Each circle represents the supernatant p24 level in an individual well seeded with 2X10^5^ resting CD4^+^ T cells isolated from pre-ATI leukapheresis samples. Maximum likelihood IUPM estimates were provided for mQVOAs (see Methods). (d) Percent reduction in infectious units per million cells (IUPM) relative to control IgG across standard progressors (SPs), illustrating wide functional heterogeneity in aNAbs against inducible, infectious reservoir virus; open circles represent values from a previously described cohort^28^. SP206 lacked detectable HIV-specific antibodies due to being treated during hyperacute infection (HAI) and served as a control for study rebound dynamics in the absence of neutralizing antibody responses. Bars indicate median with IQR. (e) Time to rebound was measured as the time in days to ≥200 HIV-1 RNA copies/mL, and was variable across all study participants. The LOD for the VL measurements was 30 HIV-1 RNA copies/mL. (f) Among SPs with measurable viral outgrowth, those with higher aNAb-mediated suppression of outgrowth showed significantly delayed rebound by Kaplan-Meier analysis (p = 0.0025, log-rank test). (g) Plasma HIV-1 RNA rebound trajectories for representative phenotypes (SP with HAI, SP, VC, EC), modeled by exponential growth curves. (h) Correlation of pre-ATI IgG-mediated suppression of ex vivo outgrowth with *in vivo* plasma viral load exponential growth rate during the ATI for SPs, including the HAI participant who lacked HIV-1 specific antibodies. Higher *ex vivo* suppression of outgrowth by aNAbs correlates with slower viral load exponential growth rates (p=0.0409, simple linear regression). (i) Plasma HIV-1 RNA doubling time during the ATI for representative phenotypes (HAI, SP, VC, EC), modeled by exponential growth curves. (j) Correlation of pre-ATI IgG-mediated suppression of *ex vivo* outgrowth with *in vivo* plasma viral load doubling time during the ATI for SPs, including the HAI participant. Higher *ex vivo* suppression of outgrowth by aNAbs correlates with delayed viral load doubling time during the ATI (p=0.0060, simple linear regression).

We next examined whether greater suppression of viral outgrowth by pre-ATI autologous IgG correlated with delayed viral rebound *in vivo*. As expected, time to viral rebound (defined as the interval from ART interruption to ≥200 HIV-1 RNA copies/mL) varied across participants, (range 7-81 days; median=19 days; Fig. 3e), consistent with previous studies^4,7,8^. We hypothesized that aNAbs capable of more effectively suppressing reactivation of inducible, replication-competent proviruses *ex vivo* could be associated with delayed viral rebound following ART interruption. To test this, we correlated the degree of *ex vivo* suppression by aNAbs, quantified as the percent reduction in IUPM with time to rebound. Stratification of SPs into tertiles based on IUPM reduction revealed a clear relationship between aNAb-mediated suppression and rebound kinetics: participants with no to minimal suppression by aNAbs (0-5%) rebounded earliest (median=10 days), those with intermediate suppression by aNAbs (5-34%) rebounded later (median=13 days), and those with high suppression by aNAbs (34-95.4%) experienced the longest delays (median=22 days; log-rank p test= 0.0025; Fig. 3f). Furthermore, linear regression analysis also showed a significant correlation (p=0.0170, Extended Data Fig. 5a) Thus, greater suppression of viral outgrowth by pre-ATI aNAbs *ex vivo* correlated with longer time to viral rebound *in vivo* among SPs with chronic HIV-1 infection.

We next evaluated whether the observed suppression of outgrowth was correlated with rebound kinetics *in vivo*. Plasma HIV-1 RNA was measured three times per week during ATI to capture early rebound dynamics with high resolution. Viral rebound trajectories were fit to an exponential growth model, log(Y) = log(Y_0_) + k·X, where Y is plasma HIV-1 RNA, X is days from first detectable viremia to peak, and k is the exponential growth constant; doubling times were calculated as ln(2)/k. Rebound kinetics varied by phenotype: the HAI participant treated within 10 days of HIV-1 acquisition lacked HIV-1-specific antibodies demonstrated one of the fastest viral load exponential growth rates whereas SPs exhibited relatively fast to intermediate viral load exponential growth rates, and controllers had the slowest viral load exponential growth rates among clinical phenotypes (Fig. 3g). Viral load exponential growth rates of SPs were inversely correlated with percent reduction in IUPM by aNAbs (p = 0.0409; Fig. 3h). As expected, doubling times showed the opposite trends (Fig. 3i) and positively correlated with aNAb-mediated suppression of *ex vivo* viral outgrowth (p = 0.0060, Fig. 3j). These data show that variability in rebound dynamics can be explained, at least in part, by the efficacy of autologous IgG at the time of rebound.

Finally, we assessed whether conventional reservoir and clinical parameters could explain variability in rebound dynamics in this cohort. We tested a wide range of metrics including intact and total HIV DNA by IPDA, IUPM by outgrowth assays, and other clinical parameters such as CD4^+^ T-cell nadir and duration on suppressive ART, most of which showed no significant association with time to rebound (Extended Data Fig. 5b-k). The only metric with a significant relationship to time to rebound was greater median pairwise *env* diversity within the latent reservoir, which was modestly associated with shorter time to rebound (p = 0.0121), suggesting that reservoirs harboring more genetically diverse *env* lineages contain a broader repertoire of immune escape variants capable of reactivating following ART interruption. Collectively, these findings indicate that while conventional reservoir and clinical metrics capture aspects of proviral burden, they do not reflect the selective immune pressures shaping the inducible, replication-competent reservoir and therefore fail to explain the heterogeneity in rebound timing observed across individuals.

### Waning aNAbs on ART allow some archived variants to regain rebound competency

We next examined whether waning autologous neutralizing antibody (aNAb) activity during long-term ART enables previously sensitive reservoir variants to gain rebound potential. We generated pseudoviruses carrying proviral, outgrowth, or rebound Envs and tested them against autologous IgG collected from longitudinal pre-ART and on-ART timepoints. Most participants were on prolonged ART prior to the ATI (median=9.5 years on ART; mean=10.6 years on ART, see Extended Data Fig. 1). Identical pseudoviruses were evaluated against IgG from multiple timepoints; therefore differences in inhibitory potential reflects temporal changes in aNAb concentration or abundance in plasma rather than viral evolution.

The IgG sampling timeline for a representative participant, VC102, is shown in Fig. 4a. An outgrowth variant from this participant, 102.QVA.2, was potently neutralized by all pre-ART IgG timepoints (Fig. 4b). Autologous IgG purified 5 weeks before ART initiation was highly active (*IP*_*10mg/mL*_ = 4.15) and activity improved slightly (*IP*_*10mg/mL*_ = 4.34) by 2 months on ART. Thereafter, a steady decline in aNAb potency was observed across IgG purified from 4 months to 10.6 years on ART. By 10.6 years on ART (10 weeks before ATI), inhibition decreased to *IP*_*10mg/mL*_ = 1.79, representing an approximately 230-fold reduction relative to the most potent pre-ART timepoint. Importantly, this gain in functional resistance reflects a decline in aNAb concentration over time on ART rather than the acquisition of aNAb-escape mutations.

**Figure 4.**
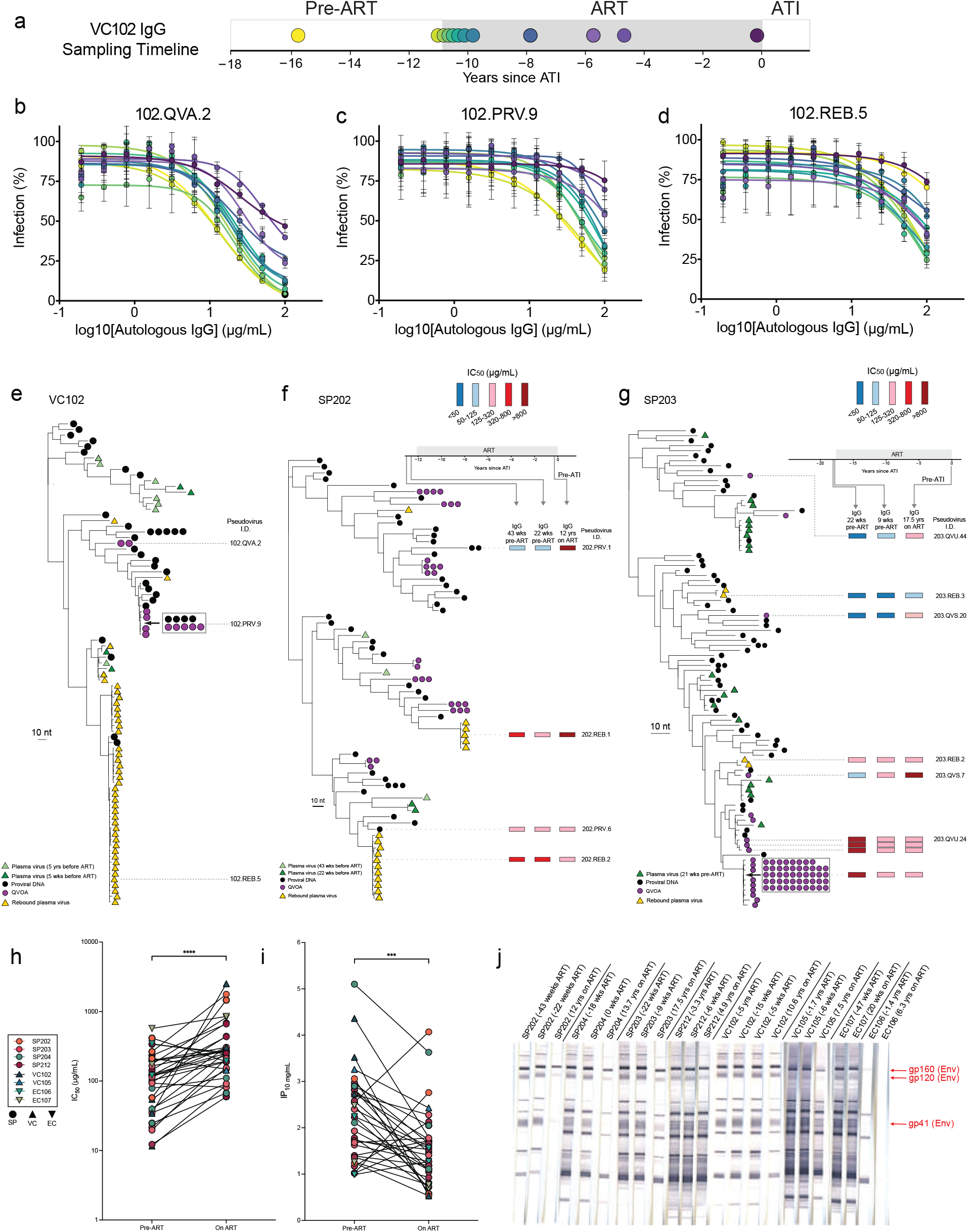
Waning autologous antibody titers during ART expand the rebound-competent HIV-1 reservoir. (a) Schematic of the sampling timeline and autologous IgG purification strategy for VC102. (b-d) Longitudinal autologous neutralization of an outgrowth variant (102.QVA.2), a reservoir variant (102.PRV.9), and a rebound isolate (102.REB.5). Neutralization potency was high at pre-ART timepoints, transiently increased after ART initiation, then declined progressively during long-term ART, resulting in complete loss of measurable neutralization for 102.PRV.9 and 102.REB.5 variants by 10.6 years on ART. (e) Maximum-likelihood *env* phylogeny illustrating genetic relationships among VC102 proviral, outgrowth, and rebound sequences. (f-g) Representative longitudinal neutralization profiles for SP202 and SP203 showing similar declines in autologous antibody potency during extended ART, leading to reduced or absent neutralization of reservoir and rebound variants. Horizontal timelines indicate autologous IgG sampling, with arrows denoting timepoints used for each neutralization assay result in relation to time on ART; continuous suppressive ART before the ATI is indicated by the dark gray shaded region on the horizontal timeline. Pseudovirus IDs are displayed to the right of the neutralization heatmaps (blue-to-red color scheme representing high-to-low aNAb potency). Viral isolates with distinct nucleotide sequences but 100% identical amino acid sequences share identical *IC*_*50*_ values, reflecting equivalent neutralization phenotypes. (h) Cohort-level comparisons of IC_50_ values of isolates tested against pre-ART versus on-ART autologous IgG (log-transformed) (p<0.0001, Wilcoxon test). (i) Cohort-level comparisons of *IP*_*10mg/mL*_ values of isolates tested against pre-ART versus on-ART autologous IgG (p = 0.0006, Wilcoxon test). (j) Western blot analysis showing qualitative reductions in gp41-, gp120-, and gp160-binding activity between purified IgG from pre-ART and on ART IgG timepoints. Negative values denoted above the western blot strips indicate the time (in weeks or years) prior to ART initiation, and the study participants remained durably suppressed on ART leading up to their participation in the REBOUND ATI trial.

Pseudovirus 102.PRV.9, carrying a proviral Env sequence, could be neutralized by autologous IgG purified prior to ART initiation (Fig. 4c). The most potent pre-ART inhibition occurred 5 weeks before ART initiation (*IP*_*10mg/mL*_ = 3.52). Thereafter, a gradual decline in aNAb potency was observed over time on ART. Autologous IgG purified at 10.6 years on ART poorly inhibited this isolate (*IP*_*10mg/mL*_ = 0.53), representing an approximately 1,000-fold decrease in neutralizing activity.

Pseudovirus 102.REB.5, carrying a representative rebound Env sequence, was poorly inhibited by IgG purified 5 years before ART (*IP*_*10mg/mL*_ = 0.78, Fig 4D). Subsequently, measurable neutralizing activity developed, as seen at 15 weeks and 5 weeks before ART initiation, the latter representing the most potent pre-ART timepoint (*IP*_*10mg/mL*_ = 2.73). Inhibition increased modestly at 1 month on ART (*IP*_*10mg/mL*_ = 4.03) but then declined progressively over time on ART. Autologous IgG purified at 10.6 years on ART had an *IP*_*10mg/mL*_ value of 1.75, representing a ∼10-fold decrease in aNAb potency relative to the most potent pre-ART timepoint. The *env* phylogeny for VC102 (Fig. 4e) provides genetic context for these variants.

Similar longitudinal patterns were observed in SP202 and SP203. In SP202 (Fig. 4f), the reservoir variant 202.PRV.1 carrying a proviral Env sequence was modestly inhibited by IgG from 43 and 22 weeks before ART initiation, with the strongest pre-ART activity corresponding to an *IP*_*10mg/mL*_ of 3.06. After 12 years on ART, inhibition declined to *IP*_*10mg/mL*_ = 0.58, representing a ∼300-fold reduction in inhibitory activity over extended time on ART. Similar trends are observed for pseudovirus 202.REB.1 (Fig. 4f).

For SP203 (Fig. 4g), pseudovirus 203.QVS.20, carrying an outgrowth virus Env sequence, aNAbs caused substantial inhibition 22 weeks before ART (*IC*_*50*_ = 29.0 µg/mL, *IP*_*10mg/mL*_ =2.82). After 17.5 years on ART, the *IP*_*10mg/mL*_ value decreased to 1.07 representing a 56-fold reduction in aNAb potency. Other pseudoviruses from SP203 (203.QVU.44, 203.QVS.7) show similar decay in aNAb potency over time on ART compared to pre-ART IgG (Fig. 4g). Similarly, a pseudovirus carrying a rebound Env sequence (203.REB.3), was neutralized by pre-ART IgG (*IC*_*50*_ = 39.5 µg/mL, *IP*_*10mg/mL*_ =2.40), but after 17.5 years on ART, the *IP*_*10mg/mL*_ value decreased to 1.59, representing a 6.5-fold decrease in aNAb potency which increased its rebound potential. These data demonstrate that rebound can arise from lineages that were not well neutralized as well as from variants that were potently inhibited by aNAbs prior to ART but became functionally resistant as aNAb concentrations in plasma waned during extended time on ART.

Analysis of 33 independent isolates from 8 study participants with paired pre-ART and on-ART IgG samples showed that aNAb potency against unique pseudoviruses carrying reservoir or rebound-derived Envs declined markedly after extended time on ART (Fig. 4h-i). The median *IC*_*50*_ value for pseudoviruses tested against the most potent pre-ART IgG sample was 121 µg/mL (IQR: 40.1-197). The median *IC*_*50*_ value for pseudoviruses tested against on-ART IgG was 258 µg/mL (IQR: 151-437), representing a significant reduction in neutralizing potency by on-ART IgG (Fig. 4h, p < 0.0001 by Wilcoxon test). Correspondingly, the median *IP*_*10mg/mL*_ value for these pseudoviruses tested against the most potent pre-ART IgG sample was 2.23 log_10_ units (IQR: 1.40-2.82). The median *IP*_*10mg/mL*_ value of pseudoviruses tested against on-ART IgG was 1.47 log_10_ units (IQR: 0.99-1.79), corresponding to a significant decline in aNAb potency on ART by 0.76 log_10_ units, or a ∼5.8-fold reduction (Fig. 4i, p = 0.0006 by Wilcoxon test). Qualitative western blot analysis (Fig. 4j) showed parallel decreases in gp41-, gp120, and gp160-binding IgG in on-ART IgG, compared to pre-ART IgG, consistent with contraction of Env-specific antibody responses after ART.

Together, these data show that pre-ART aNAb responses can mediate up to >1,000-fold inhibition, depending on the variant, but neutralization activity declines substantially due to waning aNAb concentrations in plasma following ART initiation. Thus, as humoral immune pressure shifts over time, this process can govern which variants may persist as rebound-competent lineages or can regain rebound potential as aNAb pressure wanes.

### Rebound originates directly from archived reservoir variants, not recombination

To determine whether rebound viruses originate from latently infected resting CD4^+^ T cells, we compared *env* sequences from rebound plasma viruses with proviral DNA and viral outgrowth (QVOA) isolates derived from purified resting CD4^+^ T cells from peripheral blood collected a median of 9 weeks prior to the ATI. Maximum-likelihood phylogenetic trees showed that rebound viruses were often similar to existing reservoir variants rather than derived from new, divergent lineages. Representative *env* phylogenies for SP211 and SP213 (Fig. 5a-b) illustrate rebound viruses clustering as exact matches to latent proviruses and aNAb-resistant proviruses detected in large clones of infected cells using the QVOA. Similar topologies were observed in other participants, with rebound isolates genetically linked to large, aNAb-resistant clones within the latent reservoir (see Fig. 2g-h and Fig. 4e-g). In contrast, large, aNAb-sensitive clones did not contribute to rebound (see Fig. 2h-i). Together, these findings demonstrate that rebound viruses can be exact or near-exact matches to archived proviruses in circulating resting CD4^+^ T cells.

**Figure 5.**
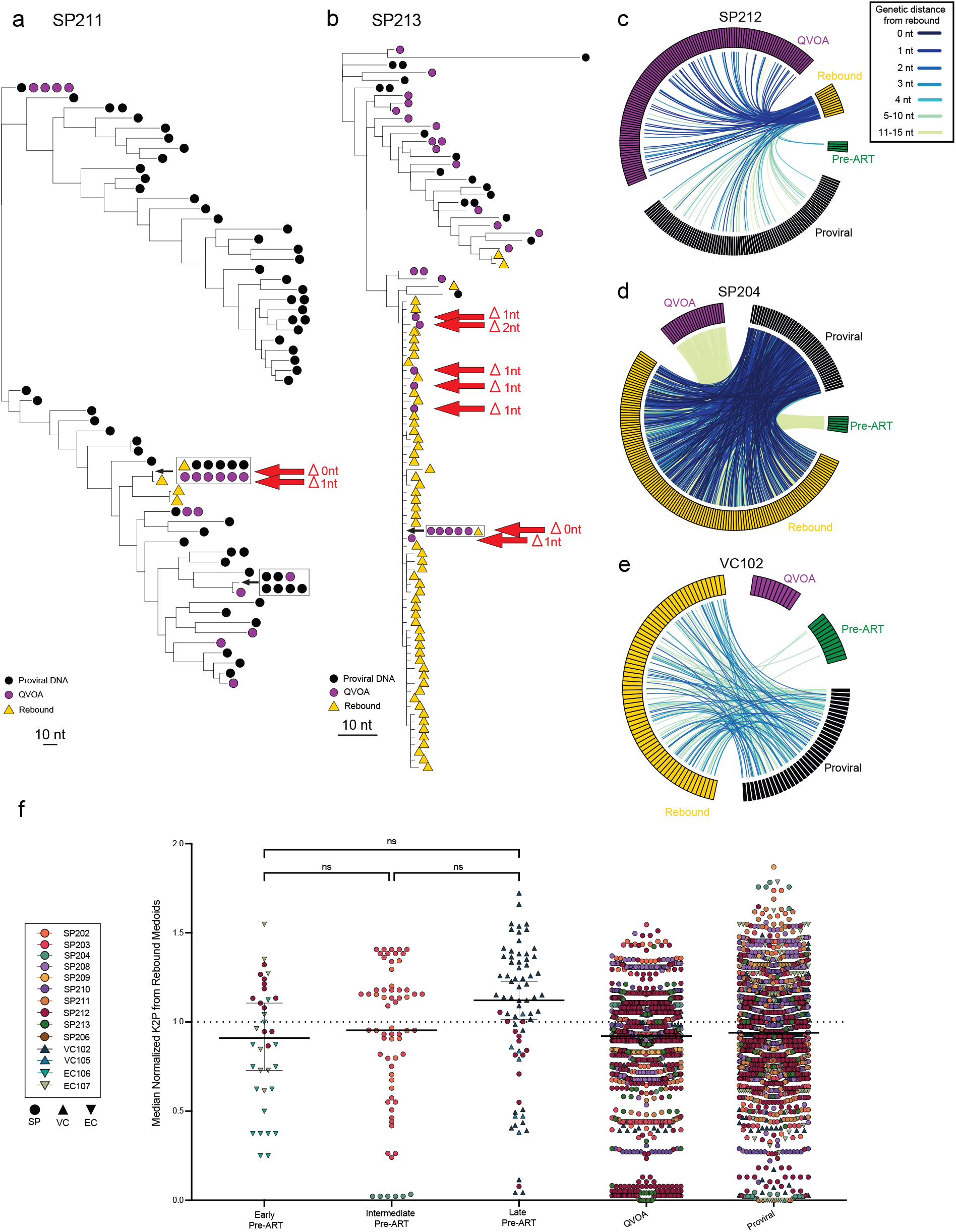
Rebound originates from archived reservoir variants as opposed to recombination. (a-b) Representative maximum-likelihood, midpoint rooted *env* phylogenies (bootstrap replicates: n=1,000) for study participants (a) SP211 and (b) SP213 showing exact (0 nt) or near-exact (1-2 nt) matches between latent reservoir (proviral or QVOA-derived) and rebound plasma *env* sequences. These matches are indicated by red arrows, which confirm direct derivation from archived proviruses. Pre-ART plasma virus sequences (green triangles), proviral DNA (black circles), QVOA-derived isolates (purple circles), and rebound plasma viruses (yellow triangles) are represented on midpoint rooted *env* phylogenies. Representative circos plots (c-e) depict all *env* sequences per participant, with colored links connecting each rebound virus to its nearest genetic neighbor among proviral, QVOA-derived, or pre-ART plasma sequences, illustrating the relationships observed in (c) SP212, (d) SP204, and (e) VC102. (f) Normalized Kimura two-parameter distances between rebound medoids and pre-ART plasma sequences stratified by time before ART initiation (early, intermediate, late). Comparable genetic proximity across tertiles (p=ns, Kruskal-Wallis test followed by Dunn’s multiple comparisons test) indicates that rebound-competent viruses can likely be archived throughout extensive timeframes during untreated infection, rather than preferentially seeded near the time of ART initiation.

To illustrate this relationship, we used circos plots to show the genetic links between each rebound virus and its nearest archived reservoir variant (Fig. 5c-d). In participants SP212, SP204, and VC102, rebound viruses were genetically identical or near-identical (Δ0-2 nt) to proviral and QVOA-derived isolates, demonstrating a direct genetic link between rebound viruses and proviruses detected in circulating CD4^+^ T cells. Across the cohort, 9 of 13 sequenced participants showed this pattern of close genetic relatedness between rebound and reservoir variants (Δ0-15 nt; ∼0-0.5% divergence; Extended Data Fig. 6).

To determine whether unlinked rebound viruses reflected recombination or reservoir undersampling, we analyzed 1,703 unique *env* sequences from proviral DNA, QVOA, and plasma virus across 13 ATI participants using RAPR (Los Alamos National Laboratory). Sixteen putative recombinants were identified (0.9% of all sequences; Extended Data Table 3), the majority (14 of 16) being recombinants between two different archived proviruses. Only 1 of 487 rebound sequences (<0.2%) was classified as a putative recombinant between two distinct proviral *env* sequences. Thus, recombinant viruses were exceedingly rare among the initial rebounding virus population, suggesting that recombination is likely not required to drive initial viral rebound post-ART.

Among the 8 participants with available pre-ART plasma, 6 showed clear genetic relatedness between pre-ART and rebound viruses (Fig. 5c-e; Extended Data Fig. 6). To evaluate whether rebound viruses were preferentially seeded near the time of ART initiation, we analyzed 70 pre-ART *env* sequences collected 1 month to 3.3 years before ART initiation (Supplementary Fig. 1; Supplementary Table 2). Genetic divergence between pre-ART plasma viruses and rebound medoids was assessed using normalized Kimura two-parameter (K2P) distances (see Methods). In this analysis, a normalized value of 1 represents the median genetic distance between a pre-ART plasma sequence and the rebound medoid, and values <1 indicate closer-than-expected relatedness. Pre-ART sequences were stratified by proximity to ART initiation (early, 40-17 months; intermediate, 16-4 months; late, 4-1 month), and normalized distances did not differ significantly between tertiles (p=ns; Fig. 5f), indicating that rebound viruses were not preferentially derived from the most recently seeded variants in our dataset. However, this analysis is limited by sparse longitudinal pre-ART sampling from within three years of ART initiation and the inclusion of some controllers.

Together, these data demonstrate that recombination is not required for viral rebound and that rebound viruses predominantly arise through reactivation of archived proviruses with sequences directly linked to circulating resting CD4^+^ T cells sampled in the peripheral blood, although the activation of latently infected cells is likely to occur in the secondary lymphoid organs.^63–65^ Although the timing of reservoir seeding cannot be precisely determined, we did not find that rebound came exclusively from variants seeded within that last few months before ART initiation. Our findings do not support a model in which rebound is genetically linked exclusively to the most evolved variants. This interpretation is consistent with our observation that variants potently neutralized by pre-ART IgG were likely seeded earlier in untreated infection; as aNAb pressure wanes on ART, these earlier archived variants can regain rebound potential and ultimately drive viral rebound. It is also possible that some early-seeded minor variants did not elicit a potent aNAb response, allowing them to persist in the reservoir as rebound competent lineages.

## DISCUSSION

This study provides quantitative criteria defining HIV-1 rebound competency. We demonstrate that rebound following ART interruption arises from the subset of archived proviruses that are highly resistant to contemporaneous aNAbs with *IP*_*10mg/mL*_ values at the lower end of the range observed for all reservoir variants. Rebound is therefore not purely stochastic; instead, it reflects a selective process governed by contemporaneous aNAbs. The role of aNAbs in shaping HIV-1 evolution has long been appreciated, with seminal work by Richman and Shaw laboratories demonstrating that antibody pressure drives viral escape and diversification during untreated infection^30,31^. In the setting of ART interruption, Li and colleagues further showed that rebound viruses are resistant to contemporaneous autologous IgG, establishing immune escape as a defining feature of rebound^35^. Building on these foundational observations, we provide quantitative criteria for rebound competency and demonstrate that within the archived latent reservoir, a select subset of reservoir variants falling below a defined inhibitory potential threshold by aNAbs appear to give rise to initial viral rebound.

The inhibitory potential of aNAbs against reservoir viruses varied over an extremely wide range (0.4-8.2 logs of inhibition). However, of 17 rebound Env variants tested in standardized single-round pseudovirus neutralization assays, all were highly resistant to pre-ATI plasma (*IC*_*50*_>80 μg/mL, *IP*_*10mg/*mL_ ≤ 2.8). These data establish an empirical threshold of *IP*_*10mg/mL*_ values below which a variant can cause rebound and suggest that substantially higher inhibitory potential against all reservoir variants may be required to prevent rebound. For context, effective combination ART regimens that can durably suppress active viral replication to below the limit of detection have combined IP values of >5 logs.^55^ We, therefore, speculate that *IP*_*10mg/mL*_ values ≤2.77 are insufficient to prevent rebound, and a functional cure by aNAbs may require ≥5 logs of inhibition of all reservoir viruses. Further studies will be required to define this value more precisely.

These inhibitory thresholds can be compared to neutralization metrics commonly used in HIV-1 prevention and passive immunization studies^66–69^. Based on prevention trials involving passive infusion of the bNAb VRC01, Gilbert et al. proposed the predicted serum neutralization 80% inhibitory dilution titer (*PT*_*80*_) as biomarker for vaccine efficacy. A *PT*_*80*_ value > 200 was associated with 90% protection against acquisition. This means that a bNAb concentration 200 fold greater than the *in vitro IC*_*80*_ of the bNAb against a given challenge virus was required for 90% efficacy in prevention. If we assume a typical Hill coefficient of 1, this corresponds to an *IP* of 2.9 logs. Interestingly, rebound in our cohort selectively emerged from the subset of reservoir variants exhibiting comparatively weak contemporaneous aNAb-mediated inhibition (*IP* < 2.8 logs). Although prevention studies and treatment interruption studies differ substantially in biologic context and antibody composition, there appears to be remarkable similarity between threshold neutralization values required to prevent acquisition (*IP* > 2.9 logs) and rebound (*IP* > 2.8 logs).

We further demonstrated that potency of aNAbs against inducible, replication-competent reservoir virus correlated with rebound kinetics and timing. Participants whose pre-ATI IgG more effectively suppressed *ex vivo* outgrowth of their own reservoir viruses rebounded later, with slower exponential growth rates and longer doubling times. This indicates that in some cases, aNAbs exert enough selective pressure on rebounding virus to improve rebound outcomes. Conventional reservoir and clinical metrics, which do not account for immune pressure, showed no association with time to rebound. Notably, extended time on ART did not confer durable post-ART control in the REBOUND cohort, consistent with the persistence of inducible, infectious proviruses in people on long-term ART^70^. Thus, rebound timing and kinetics reflect a dynamic interplay between proviral reactivation and host antibody pressure at ART interruption, a finding that can be leveraged for studies on HIV cure.

In the context of cure, it is important to note that we also found evidence that prolonged ART progressively erodes aNAb potency. In the absence of antigenic stimulation, Env-specific antibody levels may decline. Bonsignori et al.^71^ reported anti-Env antibody half-lives of 33-81 weeks after ART suppression. We show here that aNAb activity wanes during extended ART in people treated during chronic infection. As inhibitory potential falls below the protective threshold, previously neutralized variants gain rebound competency. Boosting these responses during ART may help contain the rebound-competent reservoir. Although renewed antigen exposure during ATI could potentially boost pre-existing Env-specific memory B-cell responses, prolonged viremia during ATI might also support additional de novo aNAb maturation and evolution, similar to what occurs during untreated HIV-1 infection. In the present cohort, ART was intentionally restarted relatively early in standard progressors, limiting the duration of viremia and our ability to define these humoral dynamics more precisely. Future studies with longitudinal sampling across varying durations of ATI-associated viremia will be important for determining the extent to which rebound antigen exposure can stimulate additional aNAb maturation, whether such boosted responses subsequently wane following ART reinitiation, and how rapidly rebounding viruses acquire additional escape mutations under renewed antibody pressure. These considerations further support therapeutic strategies aimed at boosting or maintaining potent aNAb responses during suppressive ART prior to ATI, including therapeutic vaccination or sequential immunogen boosting approaches.

Importantly, the timing of ART initiation fundamentally shapes these humoral trajectories. Previous studies have shown that in individuals treated during acute infection, aNAb responses can improve during early ART against pre-ART plasma virus^35,36,72^ reflecting preserved germinal-center architecture, functional follicular dendritic cells (FDCs), and intact Tfh support. Sustained antigen exposure on FDCs can thus promote continued affinity maturation and durable persistence of aNAbs specific for the more limited number of Env variants that arise prior to ART. In contrast, chronic infection is characterized by increased reservoir size and diversity, disrupted lymphoid architecture, FDC depletion, and immune imprinting on earlier Env variants, limiting B cell renewal and gradual decay of Env-specific memory^73–77^. These distinct immunologic environments may explain why early ART initiation preserves the development of potent aNAb responses, whereas in chronically-treated individuals, aNAb potency diminishes during extended ART.

We provide evidence that rebounding viruses originate from the latent reservoir. Phylogenetic analyses reinforced the model of immune-dependent rebound competency. Maximum-likelihood trees and circos plots showed that rebound viruses were often identical or nearly identical to proviral or QVOA-derived isolates from resting CD4^+^ T cells in peripheral blood. Although it is likely that rebound is initiated by antigen-driven activation of latently infected cells in the lymphoid organs^16,63–65^, our studies show that with sufficient sampling the circulating pool of resting CD4^+^ T cells can often include variants similar or identical to the ones that cause rebound. The recovery of exact ∼3 kb *env* sequence matches between reservoir and rebound viruses, despite the extreme *env* diversification in chronically treated individuals, provides strong sequence-level evidence for a direct genetic relationship between latent proviruses and rebound viruses. Previous studies have shown that large, infected cell clones carrying replication-competent proviruses often do not give rise to rebound^24–27^. Our results are consistent with the idea that, at least in some cases, these clones can harbor aNAb-sensitive virus, as previously suggested^29^. Conversely, in some cases, large clones harboring aNAb-resistant virus are directly genetically linked to initial rebounding virus. Instances where aNAb-resistant singletons or clones did not give rise to rebound may simply reflect that these proviruses were not reactivated during the ATI. Nevertheless, reservoir sampling remains inherently incomplete, and we cannot exclude contributions from additional anatomical or cellular reservoirs not directly sampled in this study. In particular, tissue-resident CD4^+^ T cells and CNS-associated or myeloid cellular compartments have increasingly been investigated as potential contributors to HIV persistence and rebound^78–80^. Future investigations integrating intensive sampling across tissue compartments will be important for defining their relative contributions to viral rebound.

Among 13 participants, only 1 of 487 rebound viruses (<0.2%) exhibited any evidence of recombination between two distinct proviral *env* sequences, and it is possible that the recombined variant could be detected in the reservoir with additional sampling from this elite controller. Recombination, therefore, contributes negligibly to initial viral rebound following ART interruption. Previous studies have rarely detected reservoir-rebound matches^11,24–27,39^; this is likely attributed to insufficient depth of reservoir sequencing. Deeper sequencing achieved in our study revealed direct genetic links between rebound viruses and latent proviruses in circulating CD4^+^ T cells in 9 of 13 ATI participants.

Together, these genetic and functional data indicate that rebound competency is determined not simply by genetic intactness; rather it is a property shaped in real time by aNAb pressure. This dynamic relationship between reservoir persistence and humoral decay forms the basis of a moving target model of rebound competency in people who initiate ART during the chronic stage of HIV-1 infection (Extended Data Fig. 7). As antibody potency declines on ART, previously neutralized lineages can regain rebound potential. Importantly, because many neutralization assays in this study used contemporaneous on-ART IgG collected after prolonged suppressive therapy, reduced sensitivity of rebound variants cannot necessarily be attributed solely to the accumulation of fixed Env escape mutations. In some cases, variants classified as not neutralized during ATI may instead reflect waning or loss of autologous antibody specificities over time on ART, or a combination of both processes, further supporting the concept that rebound competency is dynamically shaped by contemporaneous humoral immune pressure. Thus, which archived proviruses are rebound-competent at the time of ART interruption depends on contemporaneous aNAb pressure, a principle with direct implications for ATI design and cure studies. Other arms of the immune response, including CD8^+^ T-cell responses^81,82^, type I interferons^83^, and antiviral monocytes^84^, have also been implicated in the rebound process. Our findings provide a quantitative framework based on aNAb inhibitory potential that can be extended to these additional immune pathways to holistically define the rebound-competent reservoir.

Most ATI trials use time to viral rebound as a primary clinical endpoint measurement to assess therapeutic intervention efficacy^7,8,85,86^. However, our findings suggest that this metric is intrinsically confounded by aNAbs, which exert selective immune pressure on inducible reservoir variants. Individuals with stronger pre-ATI antibody responses rebounded later and with slower viral replication kinetics, independent of standard reservoir size measurements. Consequently, delays in rebound may, in some cases, reflect autologous host immune response rather than direct intervention effects. Accounting for this immune component is essential when interpreting ATI outcomes and comparing across studies. Integrating longitudinal measurements of aNAb potency and specificity across more individuals will enable more accurate discrimination between immune-mediated versus intervention-induced delays in time to rebound.

Interventions that preserve or restore potent aNAb activity capable of achieving high *IP*_*10mg/mL*_ values against all inducible, replication-competent reservoir variants could potentially sustain HIV-1 remission even without measurable reductions in reservoir size. Mapping reservoir virus resistance to contemporaneous aNAbs will be critical for precision immunogen design, enabling vaccine antigens to elicit broad and potent responses against resistant lineages. Because composition and potency of aNAbs can shift during long-term ART, it may be helpful to dynamically tailor immunogens to each individual’s evolving humoral and reservoir landscape. In this framework, personalized mRNA vaccines encoding autologous Env sequences could reinforce antibody pressure as immunity wanes, maintaining high inhibitory potential against archived resistant variants and preventing viral rebound.

In conclusion, our data show that the inhibitory potential of aNAbs sets quantitative limits on the rebound-competent HIV-1 reservoir. As aNAbs wane, the inhibitory potential exerted on reservoir variants shifts, allowing previously neutralized lineages to regain rebound competency. These limits provide a quantitative framework for interpreting rebound outcomes and for designing interventions that induce and sustain immune responses above the rebound threshold to promote durable HIV-1 remission. Together, with recent observations demonstrating that potent aNAb responses and polyfunctional T cells can enforce long-term post-intervention control in PWH^72,87^, our results mechanistically demonstrate that aNAbs are central determinants of whether HIV-1 rebounds or remains durably suppressed after ART interruption.

## METHODS

### Study participants

Adults living with HIV on stable suppressive antiretroviral therapy (ART) were enrolled at the University of California-San Francisco (UCSF) as part of the Researching Early Biomarkers of Unsuppressed HIV Dynamics (REBOUND) ATI study (ClinicalTrials.gov NCT04359186). Eligibility criteria included documented HIV-1 positivity, sustained viral suppression (≤30 HIV-1 RNA copies/mL) for at least six months, and CD4^+^ T-cell counts sufficient to safely undergo leukapheresis. Viral rebound following supervised ART interruption was monitored according to the REBOUND ATI clinical protocol, and ART was reinitiated upon confirmed plasma HIV-1 RNA rebound that met restart criteria or at the participant’s request. Additional details are provided in the SI.

### Resting CD4^+^ T cell and plasma isolation

Peripheral blood mononuclear cells (PBMCs) and contemporaneous plasma were isolated as previously described^29^.

### Genomic DNA extraction and IPDA

Genomic DNA extraction and intact proviral DNA assay (IPDA) were conducted as previously described^88^.

### Autologous IgG purification

Plasma was heat-inactivated was and processed using NAb Protein A Plus Spin Columns (Thermo Fisher Scientific, #89956) following the manufacturer’s protocol to purify autologous IgG antibodies as previously described^29^. Antibodies were extensively dialyzed to remove residual antiretroviral drugs.

### HIV-1 Antigen-Reactive IgG Western Blot Analysis

Qualitative detection of HIV-1 antigen reactivity by purified participant IgG was performed using the GS HIV-1 Western Blot Kit (Bio-Rad Laboratories, #32508) according to the manufacturer’s protocol, and as previously described^29^

### Modified quantitative viral outgrowth assay (mQVOA)

Modified QVOAs were performed as previously described.^29^ The frequency of cells harboring inducible, replication-competent proviruses was expressed as infectious units per million (IUPM) resting CD4^+^ T cells, calculated using the maximum-likelihood method developed by Rosenbloom et al.^89^ and available at https://silicianolab.johnshopkins.edu. QVOAs will all-negative well outcomes were provided a median posterior estimate and an upper bound value at the 95^th^ percentile of the posterior distribution, as previously described^89^.

### Viral RNA isolation from mQVOA supernatant and cDNA synthesis

HIV-1 RNA was extracted from the supernatant of p24 positive QVOA wells using the Viral RNA Isolation Kit (Zymo Research, R1041) per manufacturer’s protocol, and *env*-specific complementary DNA (cDNA) was synthesized using SuperScript III Reverse Transcriptase (Thermo Fisher Scientific, 18-080-044) and the primer envB3out (5′-TTGCTACTTGTGATTGCTCCATGT-3′), and as previously detailed^29^.

### Single-genome env sequencing of outgrowth virus

HIV-1 *env* sequences were amplified from genomic DNA isolated from resting CD4^+^ T cells under limiting-dilution conditions to achieve ≤30% PCR-positive wells, as previously described^29^. For participant SP209, a custom reverse nested primer (envB3in_209, 5′-TTTGACAGCTTGCCCCCC-3′) was used to accommodate minor sequence variation in the primer binding site.

### Single-genome env sequencing from proviral DNA

Proviral HIV-1 *env* sequences were amplified from genomic DNA and sequenced as previously described^29,70^.

### cDNA synthesis and single-genome env sequencing of pre-ART plasma HIV-1

Viral RNA was extracted from pre-ART plasma and converted to cDNA for *env* gene amplification as recently described^72^.

### Single-genome env sequencing from rebound plasma virus

Rebound plasma viral sequences from samples with VL>1000 were generated using the established SMRT-UMI protocol^90^. Sequenced products were filtered with the PORPIDpipeline^90^. Samples with VL<1000 were processed via a single-genome amplification (SGA) method. Viral RNA was isolated from plasma with the QIAamp Viral RNA Mini Kit (QIA-GEN) and was used as template for cDNA synthesis without SMRT-UMI oligos. HIV cDNA was amplified at limiting dilution to ensure positive PCRs contained only a single cDNA template according to the PCR conditions listed above. PCR 2 primers contained 5’ tag sequences complementary to unique index primers added during a third PCR to allow for pooling of positive PCRs for sequencing.^91^ Sequences from both methods were aligned to HXB2, trimmed to the Env reading frame, and non-functional sequences were removed.

### Pseudovirus generation

Env expression constructs were generated using the pcDNA3.4-TOPO TA Cloning Kit (Thermo Fisher Scientific, #14697). For rebound isolates where *env* was not available as PCR amplicons, custom pcDNA3.4-TOPO vectors containing participant-specific *env* sequences were synthesized by Integrated DNA Technologies (IDT). Transfection of HEK293 cells and harvesting of pseudoviruses were conducted as previously described^29,56,72^.

### Neutralization assays and calculation of IP

TZM-bl cells were used to determine antibody neutralization potency, as previously described.^29,92^ For each isolate, the fraction of uninhibited infection (*f*_*u*_) was determined by normalizing luminescence in antibody-treated wells to the pseudovirus-only control after background subtraction. Neutralization potency was quantified by the half-maximal inhibitory concentration (*IC*_*50*_) and the instantaneous inhibitory potential (*IP*), calculated as previously described^53,54,59^. Additional method details are provided in the SI.

### Sequence and phylogenetic analyses

Consensus HIV-1 *env* sequences from pre-ART plasma, proviral DNA, outgrowth cultures, and rebound plasma were codon-aligned using GeneCutter (Los Alamos National Laboratory, https://www.hiv.lanl.gov/content/sequence/GENE_CUTTER/cutter.html) and manually inspected in BioEdit. Sequences with ≥2 stop codon(s), large deletions and/or hypermutation were filtered as part of our quality control process and excluded from downstream analyses. Duplicate sequences were removed with ElimDupes (https://www.hiv.lanl.gov/content/sequence/elimdupesv2/elimdupes.html) at 100% identity while output summary files were retained to preserve information on sequence matches and redundancy metrics. Maximum-likelihood phylogenetic trees were inferred using IQ-TREE v3.0 under the general time-reversible model with 1,000 bootstrap replicates and a gamma distribution of 4 categories. Trees were midpoint-rooted and visualized in MEGA11 before annotation in Adobe Illustrator. Pairwise genetic distances among unique sequences were calculated using the Kimura 2-parameter (K2P) model.

### Recombination analysis

Within-participant recombination among recovered HIV-1 *env* sequences was evaluated using the Recombination Analysis Program (RAPR; Los Alamos National Laboratory HIV Database, https://www.hiv.lanl.gov/content/sequence/RAP2017/rap.html).

### Power analysis

To quantify the sequencing depth of rebound plasma virus, a binomial power analysis was performed, as previously described.^52^ Detection probabilities were summarized per participant (Extended Data Table 3).

## EXTENDED METHODS

### Study participants

Adults living with HIV on stable suppressive antiretroviral therapy (ART) were enrolled at the University of California-San Francisco (UCSF) as part of the Researching Early Biomarkers of Unsuppressed HIV Dynamics (REBOUND) ATI study (ClinicalTrials.gov NCT04359186). Eligibility criteria included documented HIV-1 positivity, sustained viral suppression (≤30 HIV-1 RNA copies/mL) for at least six months, and CD4^+^ T-cell counts sufficient to safely undergo leukapheresis. All study procedures were approved by the UCSF Institutional Review Board, and all participants provided written informed consent. Coded, de-identified leukapheresis and plasma samples were transferred to the Johns Hopkins University School of Medicine (JHU SOM) for downstream analyses under a separate JHU IRB approved protocol. A total of 14 participants were included in this study based on the availability of baseline leukapheresis samples that yielded sufficient numbers of resting CD4^+^ T cells for multiple parallel assays. Viral rebound following supervised ART interruption was monitored according to the REBOUND ATI clinical protocol, and ART was reinitiated upon confirmed plasma HIV-1 RNA rebound that met restart criteria or at the participant’s request.

### Neutralization assays and calculations

TZM-bl cells were used to determine antibody neutralization potency, as previously described^29,85^. Input pseudovirus preparations were titrated to ensure infections fell within the linear range. Serial dilutions of autologous IgG (100 μg/mL - 10 mg/mL) were pre-incubated with pseudoviruses at 37°C for 90 minutes. TZM-bl cells, cultured as previously described^53,54,59^, were added along DEAE-Dextran (Sigma-Aldrich #D9885, final concentration 50 μg/mL), and plates were incubated for 48 hours at 37℃. Infection was quantified by luciferase expression using the Bright-Glo luciferase assay system (Promega, #2650). For each isolate, the fraction of uninhibited infection (f_u_) was determined by normalizing luminescence in antibody-treated wells to the pseudovirus-only control after background subtraction. Neutralization potency was quantified by the half-maximal inhibitory concentration (*IC*_*50*_) and the instantaneous inhibitory potential (*IP*), calculated as previously described^53,54,59^, to integrate both potency and slope effects into a single efficacy metric. As antibody concentration increases, the fraction of unaffected infection (f_u_) declines according to the relationship:

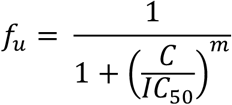

where C is the antibody concentration, and m is the slope parameter (Hill coefficient). To linearize the sigmoidal dose-response curve, we applied the median-effect equation, which describes the relationship between inhibition and antibody concentration as:

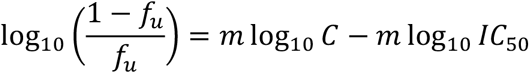

This formulation allows estimation of both *IC*_*50*_ and m from experimental data and provides a means to calculate the IP, representing the log reduction in infection events achieved at a given antibody concentration present *in vivo*:

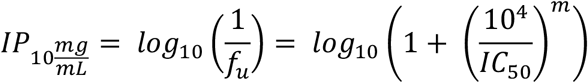

### Sequence and phylogenetic analyses

Consensus HIV-1 *env* sequences from pre-ART plasma, proviral DNA, outgrowth cultures, and rebound plasma were codon-aligned using GeneCutter (Los Alamos National Laboratory, https://www.hiv.lanl.gov/content/sequence/GENE_CUTTER/cutter.html) and manually inspected in BioEdit. Sequences with ≥2 stop codon(s), large deletions and/or hypermutation were filtered as part of our quality control process and excluded from downstream analyses. Duplicate sequences were removed with ElimDupes (https://www.hiv.lanl.gov/content/sequence/elimdupesv2/elimdupes.html) at 100% identity while output summary files were retained to preserve information on sequence matches and redundancy metrics. Maximum-likelihood phylogenetic trees were inferred using IQ-TREE v3.0 under the general time-reversible model with 1,000 bootstrap replicates and a gamma distribution of 4 categories. Trees were midpoint-rooted and visualized in MEGA11 before annotation in Adobe Illustrator.

Pairwise genetic distances among unique sequences were calculated using the Kimura 2-parameter (K2P) model, which corrects for multiple substitutions at the same site and accounts for transition-transversion bias, features that make it a more accurate estimator of evolutionary distance than p-distance, particularly for highly variable regions such as HIV-1 *env*. The K2P model minimizes underestimation of divergence that occurs when only raw nucleotide differences (p-distance) are considered. Distances were compared across compartments (proviral, QVOA-derived, and rebound plasma virus) to assess genetic relationships between latent proviruses and rebounding virus. Rebound medoids were defined as the sequences within each rebound cluster minimizing the mean pairwise K2P distance to all other sequences within the rebound cluster, computed using cophenetic distance matrices from midpoint-rooted *env* phylogenetic trees.

### Power analysis

To quantify the sequencing depth of rebound plasma virus, a binomial power analysis was performed, as previously described.^52^ The probability of detecting at least one viral lineage present at frequency p within the rebound population was calculated as: *P* = 1 − (1 − *p*)^*n*^ where n is the number of rebound *env* sequences recovered for each participant. Lineage frequencies of 10%, 15%, 25%, 30%, and 50% were examined. Participants with ≥20 rebound sequences achieved ≥90% probability of detecting all lineages comprising ≥10% of the rebound population. Participants with ≤12 sequences retained ≥95% probability of detecting dominant lineages ≥25-50% frequency. Detection probabilities were summarized per participant (Extended Data Table 3).

## EXTENDED DATA FIGURES

**Extended Data Figure 1.**
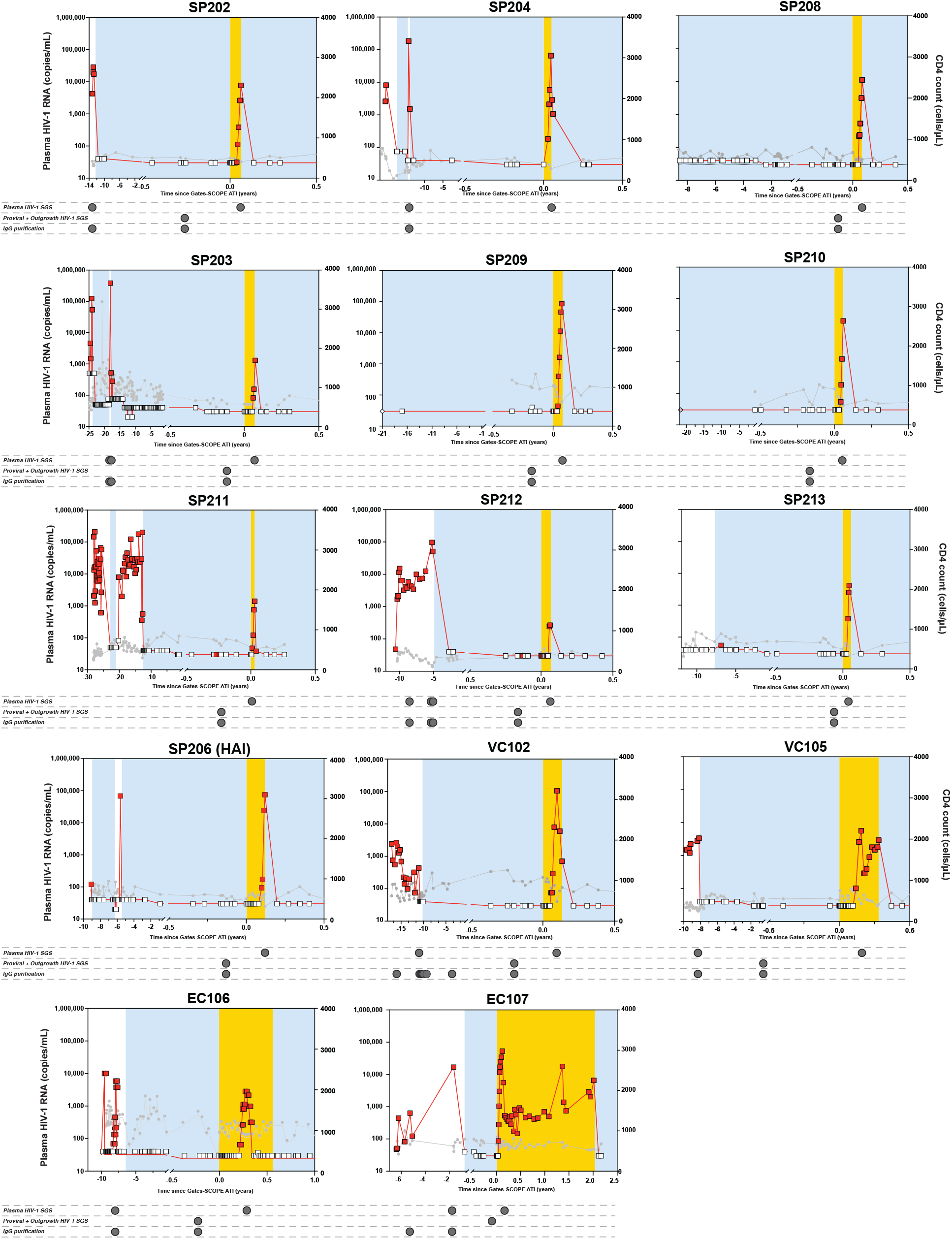
Longitudinal viral load, CD4^+^ T-cell counts, and sampling overview for all study participants. Individual participant trajectories of plasma HIV-1 RNA (left y-axis, red) and peripheral CD4^+^ T cell counts (right y-axis, gray) are shown from pre-ART baseline through long-term suppressive ART and analytic treatment interruption (ATI). Blue shaded regions denote periods of continuous ART, and yellow shaded regions indicate ATI phases during which plasma viremia was monitored three times per week to capture rebound kinetics with high temporal resolution. Horizontal tracks below each plot depict the timing and source of sample collection, including pre-ART plasma, proviral DNA from resting CD4^+^ T cells, quantitative viral outgrowth assay (QVOA) cultures, and rebound plasma isolates, as indicated by color-coded markers. Dashed vertical lines mark the start and end of ART or ATI periods. Open diamonds indicate the participant-reported date when viral suppression was first achieved if clinical documentation was unavailable. Together, these longitudinal clinical profiles illustrate the diverse treatment histories and rebound kinetics among the 14 participants, providing temporal context for all downstream sequencing, neutralization, and reservoir analyses.

**Extended Data Figure 2.**
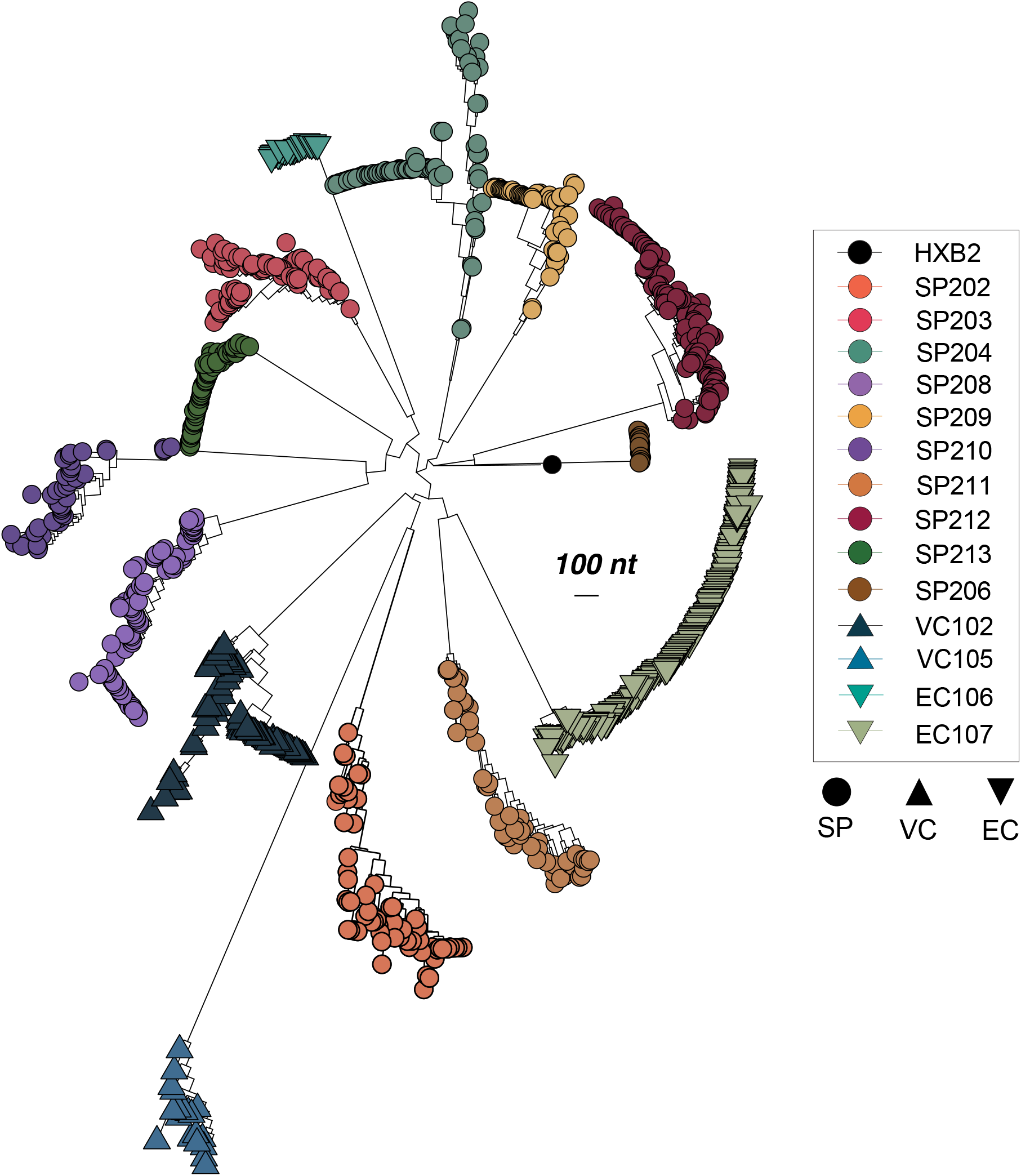
A global phylogenetic tree was inferred from all unique *env* nucleotide sequences obtained from pre-ART plasma, proviral DNA, quantitative viral outgrowth assay (QVOA) cultures, and rebound plasma across 14 ATI participants. Sequences were codon-aligned to the HXB2 reference, manually inspected to remove hypermutated, truncated, or defective sequences, and deduplicated using ElimDupes at 100% identity. Phylogenetic inference was performed using IQ-TREE v3.0.1 under the GTR+I+G4 substitution model with 1,000 ultrafast bootstrap replicates and SH-aLRT branch support. The resulting unrooted tree was midpoint-rooted and visualized in MEGA11. Branches are colored by participant, with bootstrap support values indicated for major nodes. The tree reveals distinct participant-specific monophyletic clusters with no inter-participant mixing, consistent with independent viral evolution and variable within-participant *env* diversity reflective of reservoir complexity and infection history.

**Extended Data Figure 3.**
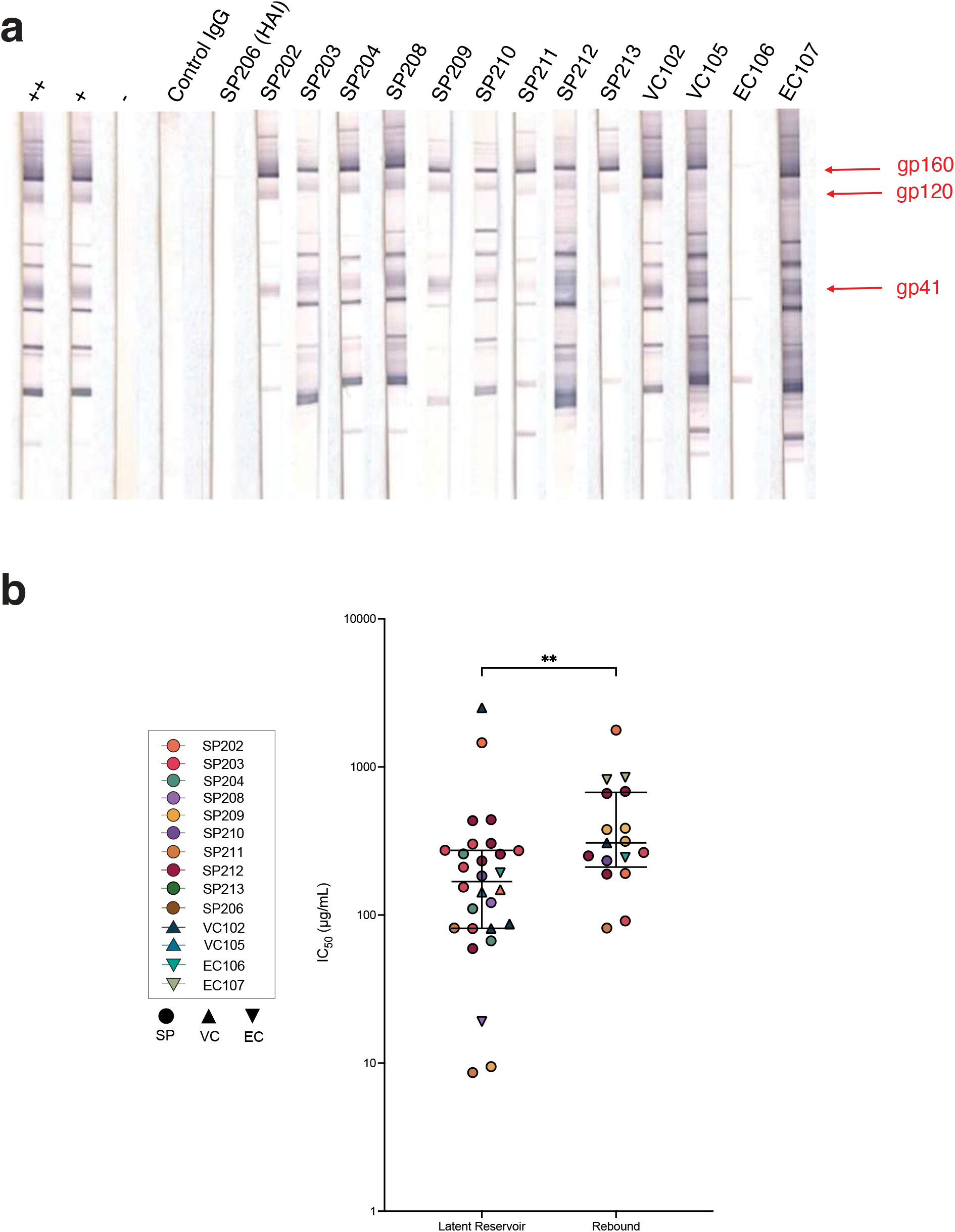
(a) Western blots assessing antibody reactivity to HIV-1 Env proteins (gp41, gp120, and gp160) in plasma from all REBOUND participants at pre-ATI timepoints. Red arrows indicate the positions of gp41-, gp120-, and gp160-reactive bands, to assess HIV-1 Env-specific IgG binding. Qualitative western blots were performed using the GS HIV-1 Western Blot Kit, according to the manufacturer’s protocol. Each blot includes Bio-Rad reference controls: **++** (high positive), **+** (low positive), and **-** (negative) controls, used to verify assay performance. Control IgG represents pooled polyclonal IgG purified from HIV-seronegative donors, included to confirm assay specificity. All blots were processed using the same exposure. (b) Comparison of *IC*_*50*_ values between reservoir viruses from Fig. 2c and rebound viruses from Fig. 2d, showing only data points from the REBOUND cohort. Values for reservoir pseudoviruses from the REBOUND cohort are on average lower than *IC*_*50*_ values for the rebound isolates. The difference between the distributions of reservoir and rebound *IC*_*50*_ values in the REBOUND cohort was significant as assessed by Mann-Whitney test (p = 0.0095). Bars indicate the median and IQR.

**Extended Data Figure 4.**
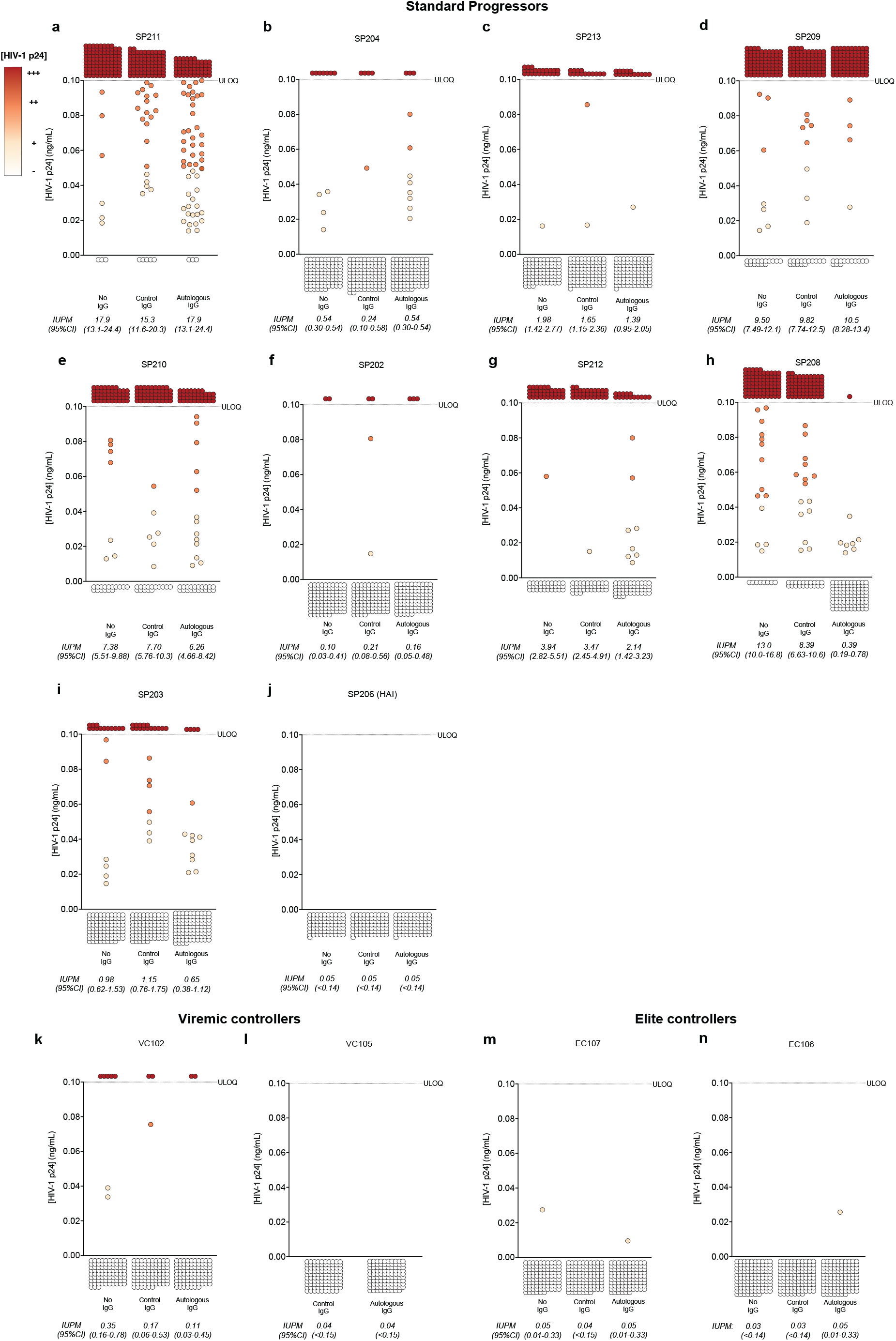
mQVOAs were performed using purified resting CD4^+^ T cells isolated from pre-ATI leukapheresis samples to assess aNAb-mediated suppression of inducible, replication-competent reservoir virus. Culture wells were activated by PHA and irradiated feeders in the presence of no IgG, control IgG from HIV-seronegative donors (50 µg/mL), or contemporaneous autologous pre-ATI IgG (50 µg/mL). Each circle represents the HIV-1 p24 antigen concentration measured by ELISA in the supernatant of a single well seeded with 2×10^5^ resting CD4^+^ T cells. HIV-1 p24 antigen levels are color-coded from white (no detectable p24 antigen) to dark red (high p24 antigen). Replicate well outcomes were used to calculate maximum likelihood estimates of infectious units per million (IUPM) resting CD4^+^ T cells (including 95% CI) for each experimental condition and participant. Participants are stratified by clinical classification: (a-j) standard progressors (SPs), (k-l) viremic controllers (VCs), and (m-n) elite controllers (ECs), highlighting phenotype-specific differences in inducible reservoir frequency and the extent of suppression mediated by pre-ATI autologous IgG.

**Extended Data Figure 5.**
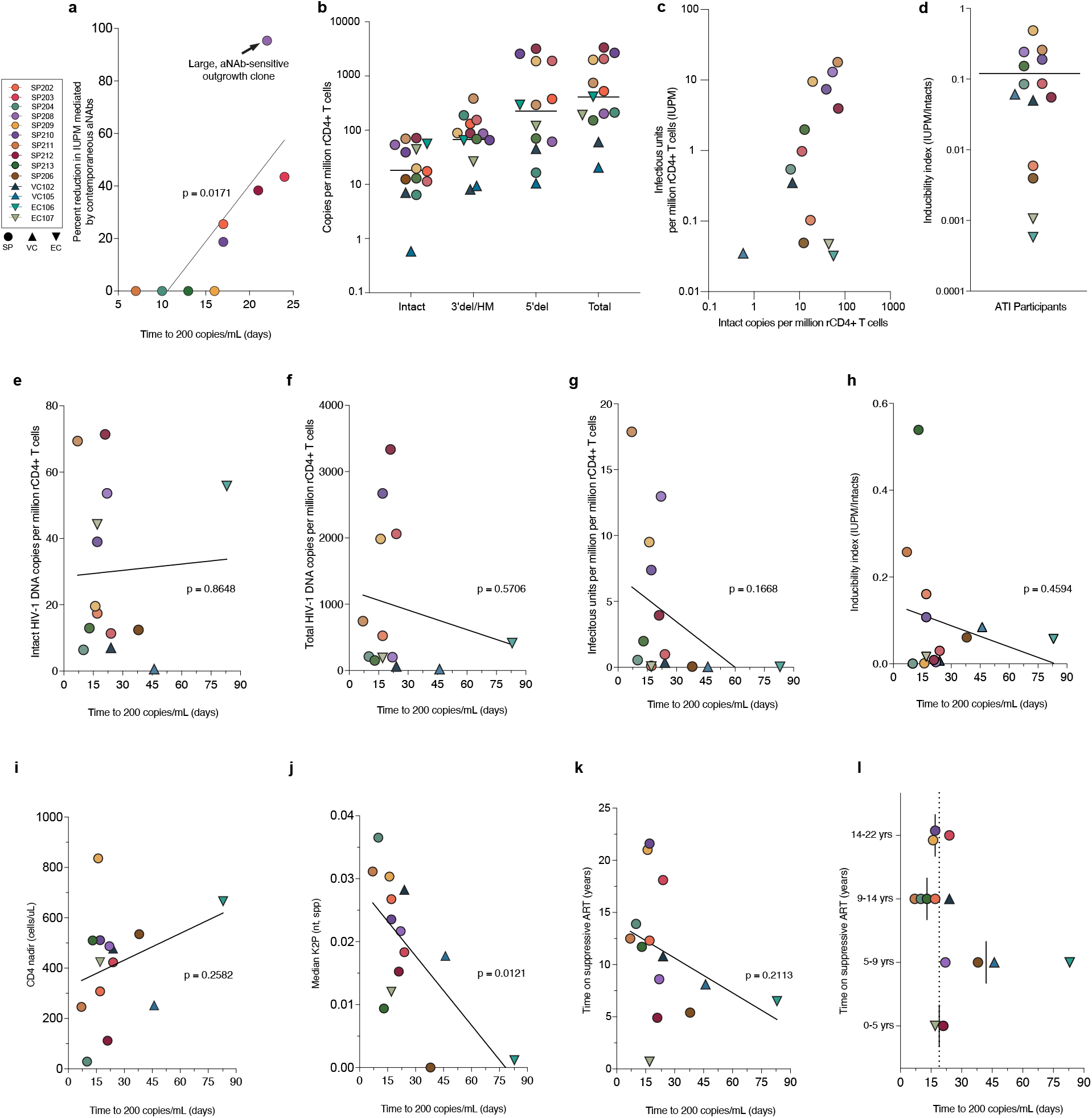
Conventional reservoir metrics and testing of correlates with time to rebound. Simple linear regression analyses between most conventional reservoir parameters and time to rebound revealed no significant correlations. (a) Linear regression of time to rebound with percent reduction by aNAbs in the mQVOA excluding VCs, ECs and HAI participants who had limited or undetectable viral outgrowth. (b) Frequency of intact, 3′ defective, and 5′ defective proviruses measured by the intact proviral DNA assay (IPDA) for all participants. (c) Relationship between infectious units per million (IUPM) resting CD4^+^ T cells, determined by quantitative viral outgrowth assay (QVOA), and the number of intact proviruses measured by IPDA. (d) Inducibility index, defined as the ratio of IUPM to intact proviral DNA copies. Correlations of time to rebound vs. (e) intact proviral DNA copies, (f) total HIV DNA copies, (g) IUPM, (h) inducibility index, (i) CD4^+^ T-cell nadir, (j) median pairwise K2P nucleotide divergence of HIV-1 *env* sequences, and (k) duration of suppressive ART, and (l) time to rebound stratified by quartiles of ART duration (0-5, 5-9, 9-14, and 14-22 years on suppressive ART).

**Extended Data Figure 6.**
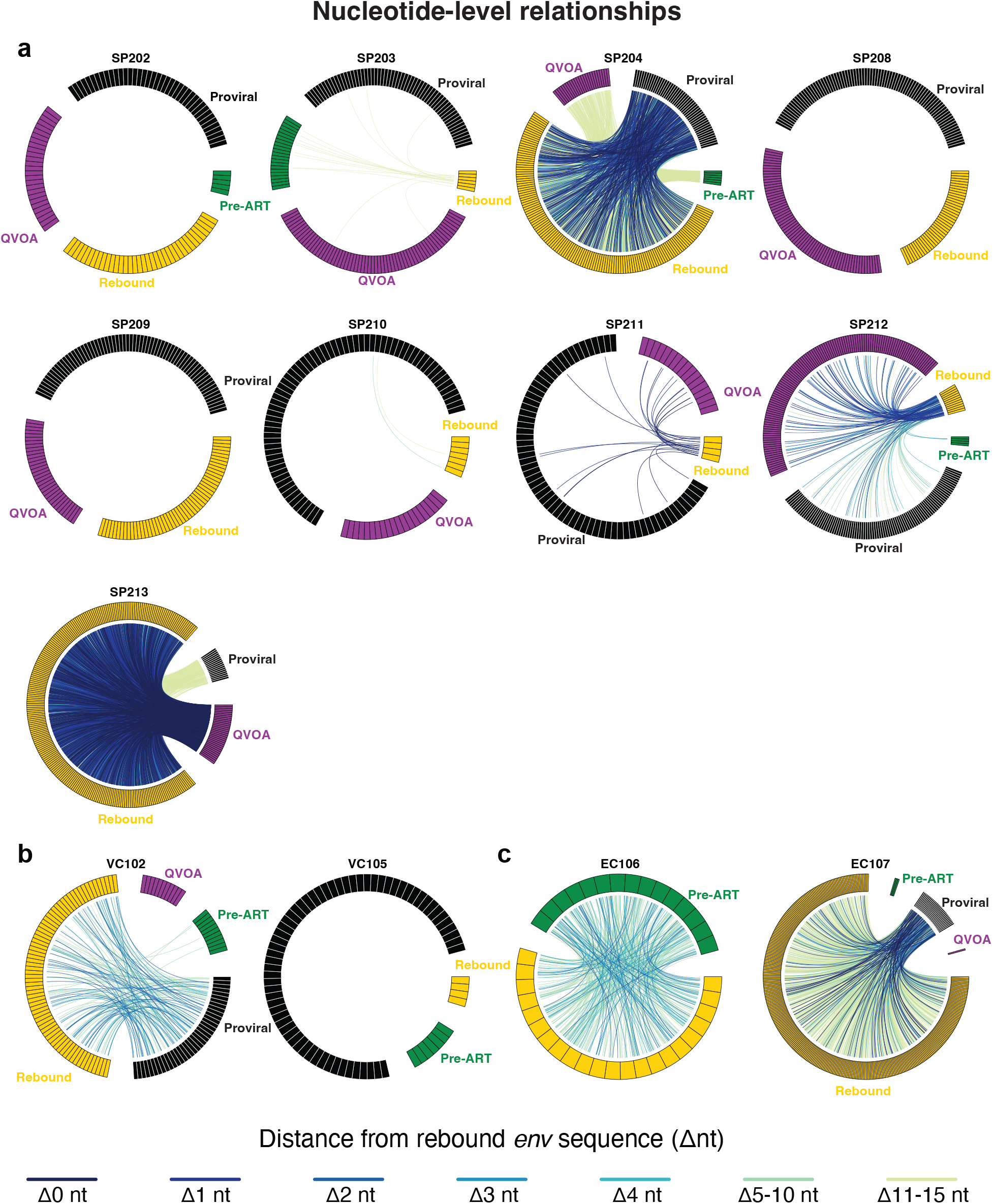
Genetic linkage between rebound and reservoir viruses across clinical phenotypes (a-c). Circos plots depict pairwise genetic relationships between rebound and reservoir-derived *env* sequences, identifying for each rebound virus its nearest archived counterpart (QVOA-derived, proviral, or pre-ART plasma) based on minimal Hamming distance. Each outer segment represents an individual *env* sequence, color-coded by source: proviral DNA (black), QVOA-derived outgrowth virus (purple), pre-ART plasma virus (green), and rebound plasma virus (yellow). Connecting arcs trace the shortest nucleotide difference between each rebound and its closest reservoir sequence. Arc color reflects genetic relatedness: dark blue links indicate exact sequence matches, with progressively greener colored links representing increasing, but still genetically similar distances between sequences (see figure key). These visualizations highlight direct genetic continuity between rebound and archived viruses, demonstrating that rebound variants frequently emerge from exact or near-identical proviral or QVOA-derived *env* sequences within the latent reservoir. (a) Standard progressors (n=9, excluding SP206 with hyperacute infection due to no recovery of reservoir sequences, (b) viremic controllers (n=2), and (c) elite controllers (n=2) are shown separately to illustrate phenotype-specific reservoir-rebound connectivity patterns.

**Extended Data Figure 7.**
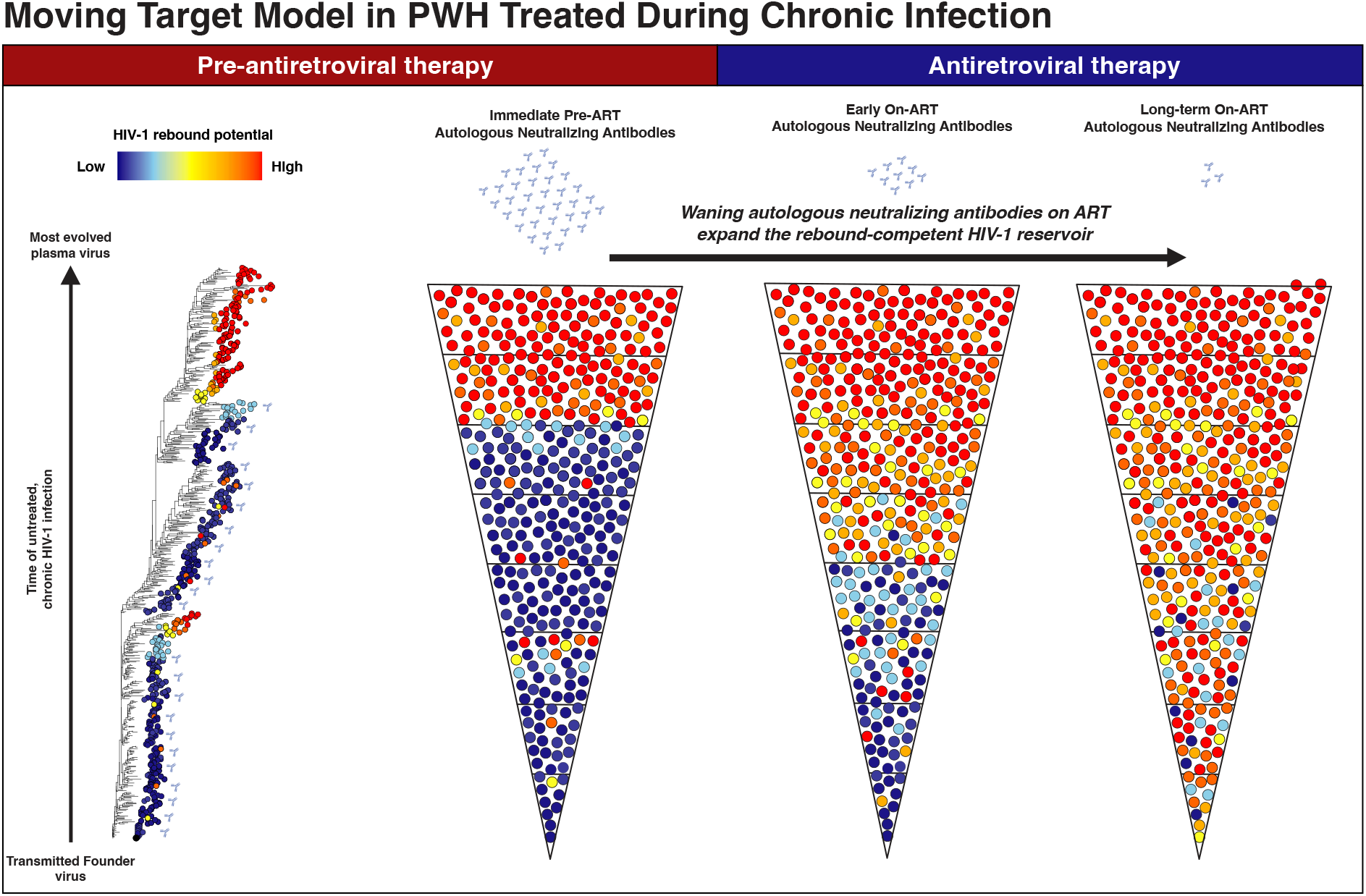
Schematic illustrating how the potency and breadth of autologous neutralizing antibodies (aNAbs) shape the rebound-competent reservoir during long-term ART. In individuals who initiated therapy during chronic HIV-1 infection, potent aNAbs initially restrict which proviruses are capable of reactivation. Over time, however, continued ART suppresses viral antigen exposure, leading to the gradual waning of aNAb titers and functional activity. As the inhibitory threshold (*IP*_*10mg/mL*_) declines, a progressively larger fraction of archived proviruses previously neutralized can regain rebound-competency. Reservoir proviruses that remain resistant to contemporaneous aNAbs at the time of ATI therefore define the functionally rebound-competent subset. Together, this “moving-target” model illustrates how the effective breadth of the rebound-competent reservoir increases over time as humoral immune pressure decays, linking antibody waning to the reactivation potential of archived proviruses.

**Extended Data Table 1.** Clinical characteristics of study participants.

| *Participant ID | Age | Race | Years since HIV-1 acquisition | Years since ART initiation | Years of documented ART | Years of estimated ART | CD4 nadir (cells/μL) | CD4 (cells/μL) | HIV-1 RNA (cps/mL) | # ART regimen |
| --- | --- | --- | --- | --- | --- | --- | --- | --- | --- | --- |
| SP202 | 41 | White | 17 | 12.3 | 11.9 | 11.9 | 308 | 379 | <30 | BIC/FTC/TAF |
| SP203 | 67 | White/European American | 31.1 | 26.5 | 17.2 | 17.2 | 423 | 578 | <30 | FTC/TAF/DTG |
| SP204 | 75 | Not specified | 36.9 | 22.1 | 4.2 | 6.3 | 29 | 457 | <30 | ABC/DTG/3TC |
| SP208 | 43 | White | 22.2 | 14.2 | 8.4 | 8.4 | 487 | 509 | <30 | FTC/TAF/DTG |
| SP209 | 63 | White/European American | 35.9 | 25.9 | 11.2 | 25.6 | 836 | 1015 | <30 | ABC/DTG/3TC |
| SP210 | 49 | Unknown | 25.2 | 21.4 | 0.3 | 21.2 | 511 | 584 | <30 | BIC/FTC/TAF |
| SP211 | 57 | White | 30.9 | 25.9 | 12.0 | 12.0 | 246 | 603 | <30 | ABC/DTG/3TC |
| SP212 | 68 | White/European American | 10.2 | 4.9 | 4.7 | 4.7 | 112 | 354 | <30 | ABC/DTG/3TC |
| SP213 | 60 | White | 13.3 | 12.3 | 6.2 | 6.2 | 510 | 551 | <30 | BIC/FTC/TAF |
| SP206 (HAI) | 60 | Mixed; White, Other | 9.0 | 9.0 | 5.4 | 5.4 | 535 | 509 | <30 | BIC/FTC/TAF |
| VC102 | 57 | White | 20.7 | 10.6 | 10.5 | 10.5 | 477 | 990 | <30 | DTG/3TC |
| VC105 | 57 | Black | 12.2 | 7.5 | 3.3 | 3.3 | 252 | 412 | <30 | ABC/DTG/3TC |
| EC107 | 32 | White | 6.1 | 1.3 | 1.2 | 1.2 | 424 | 643 | <30 | DTG/3TC |
| EC106 | 49 | White | 9.4 | 7.6 | 7.6 | 3.7 | 666 | 1472 | <30 | EVG/FTC/TAF/COBI |

*\*Participants are stratified as SP, SP (HAI), VC, or EC to indicate Standard Progressor, Standard* *Progressor with hyperacute infection, Viremic Controller, or Elite Controller, respectively.* *#ART regimens are abbreviated as follows: BIC, bictegravir; FTC, emtricitabine; TAF, tenofovir* *alafenamide; DTG, dolutegravir; ABC, abacavir; 3TC, lamivudine; EVG, elvitegravir; COBI,* *cobicistat.*

**Extended Data Table 2.** Sequencing summary and estimated detection probability for rebound plasma virus.

| * Participant ID | Proviral | QVOA | Pre-ART Plasma Virus | Rebound Plasma Virus | Total Sequences | # Estimated detection probability (%) |  |  |  |  |
| --- | --- | --- | --- | --- | --- | --- | --- | --- | --- | --- |
| | | | | | | p≥10 % | p≥15 % | p≥25 % | p≥30 % | p≥50 % |
| SP 211 | 66 | 17 | N/A | 4 | 87 | 34.4 | 47.8 | 68.4 | 76 | 93.8 |
| SP 210 | 68 | 20 | N/A | 7 | 95 | 52.2 | 67.9 | 86.7 | 91.8 | 99.2 |
| SP 202 | 40 | 27 | 5 | 36 | 108 | 97.7 | 99.7 | 100 | 100 | 100 |
| SP 203 | 60 | 65 | 22 | 6 | 153 | 46.9 | 62.3 | 82.2 | 88.2 | 98.4 |
| SP 209 | 68 | 33 | N/A | 52 | 153 | 99.6 | 100 | 100 | 100 | 100 |
| SP 208 | 85 | 69 | N/A | 41 | 195 | 98.7 | 99.9 | 100 | 100 | 100 |
| SP 204 | 48 | 25 | 6 | 140 | 219 | 100 | 100 | 100 | 100 | 100 |
| SP 212 | 94 | 121 | 4 | 12 | 231 | 71.8 | 85.8 | 96.8 | 98.6 | 100 |
| SP 213 | 17 | 32 | N/A | 238 | 287 | 100 | 100 | 100 | 100 | 100 |
| SP206 (HAI) | 0 | 0 | N/A | 136 | 136 | 100 | 100 | 100 | 100 | 100 |
| VC 105 | 63 | 0 | 7 | 4 | 74 | 34.4 | 47.8 | 68.4 | 76 | 93.8 |
| VC 102 | 37 | 11 | 11 | 69 | 128 | 99.9 | 100 | 100 | 100 | 100 |
| EC 106 | 0 | 0 | 15 | 22 | 37 | 90.2 | 97.2 | 99.8 | 100 | 100 |
| EC 107 | 30 | 1 | 3 | 323 | 357 | 100 | 100 | 100 | 100 | 100 |

\*Participants are stratified as SP, SP (HAI), VC, or EC to indicate Standard Progressor, Standard Progressor with Hyperacute infection, Viremic Controller, or Elite Controller, respectively. <sup>#</sup>*p* represents the underlying frequency of a viral lineage within the rebound plasma population; values indicate the probability of detecting at least one sequence from a lineage comprising $\geq p$ % of the total rebound virus population, calculated using the binomial framework, as previously described<sup>52</sup>.

**Extended Data Table 3.**
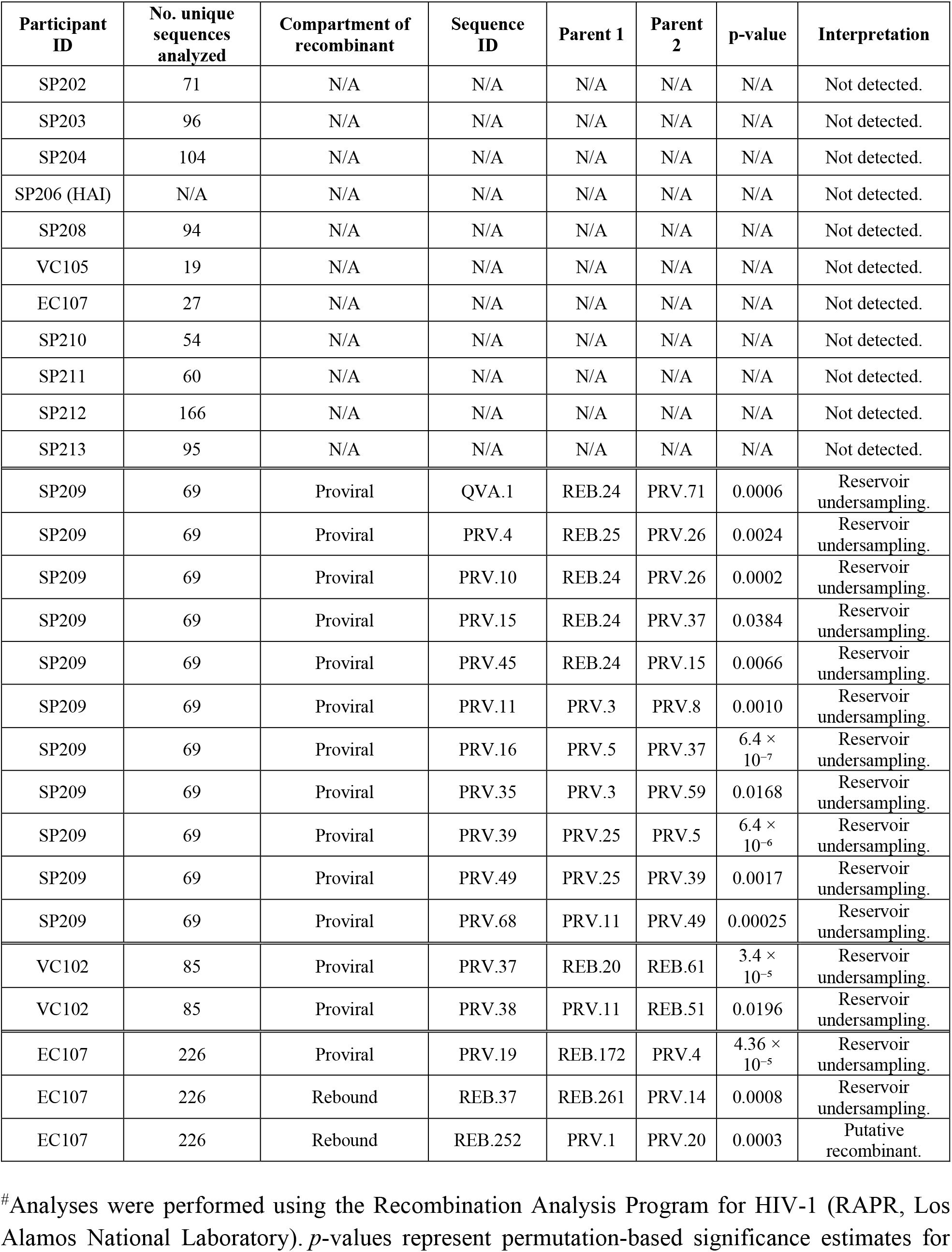
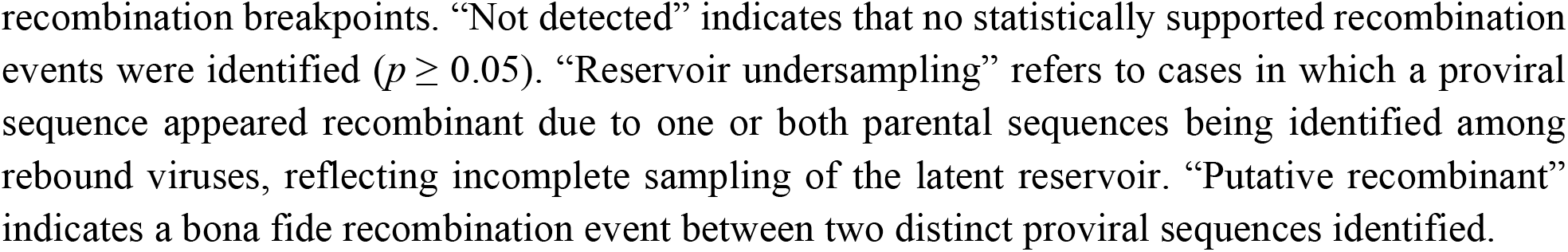
Recombination analysis of HIV-1 *env* sequences.

| Participant ID | No. unique sequences analyzed | Compartment of recombinant | Sequence ID | Parent 1 | Parent 2 | p-value | Interpretation |
| --- | --- | --- | --- | --- | --- | --- | --- |
| SP202 | 71 | N/A | N/A | N/A | N/A | N/A | Not detected. |
| SP203 | 96 | N/A | N/A | N/A | N/A | N/A | Not detected. |
| SP204 | 104 | N/A | N/A | N/A | N/A | N/A | Not detected. |
| SP206 (HAI) | N/A | N/A | N/A | N/A | N/A | N/A | Not detected. |
| SP208 | 94 | N/A | N/A | N/A | N/A | N/A | Not detected. |
| VC105 | 19 | N/A | N/A | N/A | N/A | N/A | Not detected. |
| EC107 | 27 | N/A | N/A | N/A | N/A | N/A | Not detected. |
| SP210 | 54 | N/A | N/A | N/A | N/A | N/A | Not detected. |
| SP211 | 60 | N/A | N/A | N/A | N/A | N/A | Not detected. |
| SP212 | 166 | N/A | N/A | N/A | N/A | N/A | Not detected. |
| SP213 | 95 | N/A | N/A | N/A | N/A | N/A | Not detected. |
| SP209 | 69 | Proviral | QVA.1 | REB.24 | PRV.71 | 0.0006 | Reservoir undersampling. |
| SP209 | 69 | Proviral | PRV.4 | REB.25 | PRV.26 | 0.0024 | Reservoir undersampling. |
| SP209 | 69 | Proviral | PRV.10 | REB.24 | PRV.26 | 0.0002 | Reservoir undersampling. |
| SP209 | 69 | Proviral | PRV.15 | REB.24 | PRV.37 | 0.0384 | Reservoir undersampling. |
| SP209 | 69 | Proviral | PRV.45 | REB.24 | PRV.15 | 0.0066 | Reservoir undersampling. |
| SP209 | 69 | Proviral | PRV.11 | PRV.3 | PRV.8 | 0.0010 | Reservoir undersampling. |
| SP209 | 69 | Proviral | PRV.16 | PRV.5 | PRV.37 | $6.4 \times 10^{-7}$ | Reservoir undersampling. |
| SP209 | 69 | Proviral | PRV.35 | PRV.3 | PRV.59 | 0.0168 | Reservoir undersampling. |
| SP209 | 69 | Proviral | PRV.39 | PRV.25 | PRV.5 | $6.4 \times 10^{-6}$ | Reservoir undersampling. |
| SP209 | 69 | Proviral | PRV.49 | PRV.25 | PRV.39 | 0.0017 | Reservoir undersampling. |
| SP209 | 69 | Proviral | PRV.68 | PRV.11 | PRV.49 | 0.00025 | Reservoir undersampling. |
| VC102 | 85 | Proviral | PRV.37 | REB.20 | REB.61 | $3.4 \times 10^{-5}$ | Reservoir undersampling. |
| VC102 | 85 | Proviral | PRV.38 | PRV.11 | REB.51 | 0.0196 | Reservoir undersampling. |
| EC107 | 226 | Proviral | PRV.19 | REB.172 | PRV.4 | $4.36 \times 10^{-5}$ | Reservoir undersampling. |
| EC107 | 226 | Rebound | REB.37 | REB.261 | PRV.14 | 0.0008 | Reservoir undersampling. |
| EC107 | 226 | Rebound | REB.252 | PRV.1 | PRV.20 | 0.0003 | Putative recombinant. |

